# Glucocorticoids regulate cancer cell dormancy

**DOI:** 10.1101/750406

**Authors:** Stefan Prekovic, Karianne Schuurman, Anna González Manjón, Mark Buijs, Isabel Mayayo Peralta, Max D. Wellenstein, Selçuk Yavuz, Alejandro Barrera, Kim Monkhorst, Anne Huber, Ben Morris, Cor Lieftink, Joana Silva, Balázs Győrffy, Liesbeth Hoekman, Bram van den Broek, Hans Teunissen, Timothy Reddy, William Faller, Roderick Beijersbergen, Jos Jonkers, Maarten Altelaar, Karin E. de Visser, Elzo de Wit, Rene Medema, Wilbert Zwart

## Abstract

The glucocorticoid receptor directly regulates thousands of genes across the human genome in a cell-type specific manner, governing various aspects of homeostasis. The influence of the glucocorticoid receptor is also seen in various pathologies, including cancer, where it has been linked to tumorigenesis, metastasis, apoptosis resistance, and therapy bypass. Nonetheless, the direct genetic and molecular underpinnings of glucocorticoid action in cancer remain elusive. Here, we dissected the glucocorticoid receptor signalling axis and uncovered the mechanism of glucocorticoid-mediated cancer cell dormancy. Upon glucocorticoid receptor activation cancer cells undergo quiescence, subserved by cell cycle arrest through *CDKN1C* and reprogramming of signalling orchestrated via *FOXO1*/*IRS2*. Strikingly, co-expression of these three genes, directly regulated by glucocorticoid-induced chromatin looping, correlates with a benign molecular phenotype across human cancers, whereas triple loss is associated with increased expression of proliferation/aggressiveness markers. Finally, we show that the glucocorticoid receptor signalling axis is inactivated by alterations of either the chromatin remodelling complex or *TP53 in vitro* and *in vivo*. Our results indicate that the activation of the glucocorticoid receptor leads to cancer cell dormancy, which has several implications in terms of glucocorticoid use in cancer therapy.

## Main

The Glucocorticoid Receptor (GR) is a member of the nuclear hormone receptor superfamily and a ligand-activated transcription factor^1^. This multidomain protein exerts its function through chromatin binding and communication with the transcriptional machinery, ultimately modulating expression of several hundreds of genes, ubiquitously across diverse cell types^2^. As a homeostatic regulator, GR has an imperative role in neuroendocrine integration, circadian rhythm, immune system control, and glucose metabolism^3^. GR action extends beyond general physiology as its impact can be seen in various disease types, including cancer where it has been linked to tumorigenesis^4^, metastasis^5^, apoptosis resistance^6^, and therapy bypass^7^. As demonstrated using aged mouse haploinsufficiency models, GR loss predisposes to tumour development across multiple organ systems^4^. These findings are further substantiated by studies linking glucocorticoids (GCs) to growth-arrest in cell culture based tumour models^8,9^. Despite the reported influence on tumour biology, the direct genetic and molecular underpinnings of GC-action remain elusive.

Herein, we examine the mechanism of GC-regulated cancer cell dormancy, using a multidisciplinary approach to dissect the GR axis and illustrate pathophysiological relevance thereof in human tumours. We show that GR activation leads to acquisition of quiescence, subserved by cell cycle arrest through *CDKN1C* and reprogramming of signalling orchestrated via *FOXO1/IRS2*. Strikingly, co-expression of these three genes, directly regulated by GC-induced chromatin looping, correlates with a benign molecular phenotype across human cancers, whereas triple loss is associated with increased expression of proliferation/aggressiveness markers. Finally, we demonstrate that the GR signalling axis may be inactivated either through chromatin remodelling complex (SWI/SNF) or *TP53* alterations *in vitro and in vivo*.

### Glucocorticoid receptor activation leads to cancer cell dormancy

Active GR signalling is involved in lung development^10^, differentiation^11^, and tumorigenesis^4^. Therefore, we sought to explore the molecular mechanism underlying GR action in context of lung cancer. Firstly, we analysed steroid hormone receptor expression in lung cancer patients on mRNA (110 normal tissue and 1017 cancer samples) and protein level (193 samples for GR/AR and 47 samples for ER/PR). Overall, GR was the only steroid hormone receptor expressed both on mRNA (Fig. 1a; *NR3C1*) and protein level (Fig. 1b and S1a) in most tumour samples (92,1%). Staining intensity was scored for GR immunohistochemistry, with the majority of tumours showing high intensity (3; 45,5%), while medium/low intensities were less abundant (2 and 1; 25,9 and 20,7%, respectively) (Fig. 1b). GR-restricted expression in lung cancer limits the potential of a cross-talk between steroid hormone receptors, known to modulate their function^12^. Importantly, lung cancer patients with high GR mRNA levels had a significantly higher survival probability (median survival of 111,6 months) than those with intermediate-to-low levels (median survival of 52 months) (Fig. 1c).

**Figure 1.**
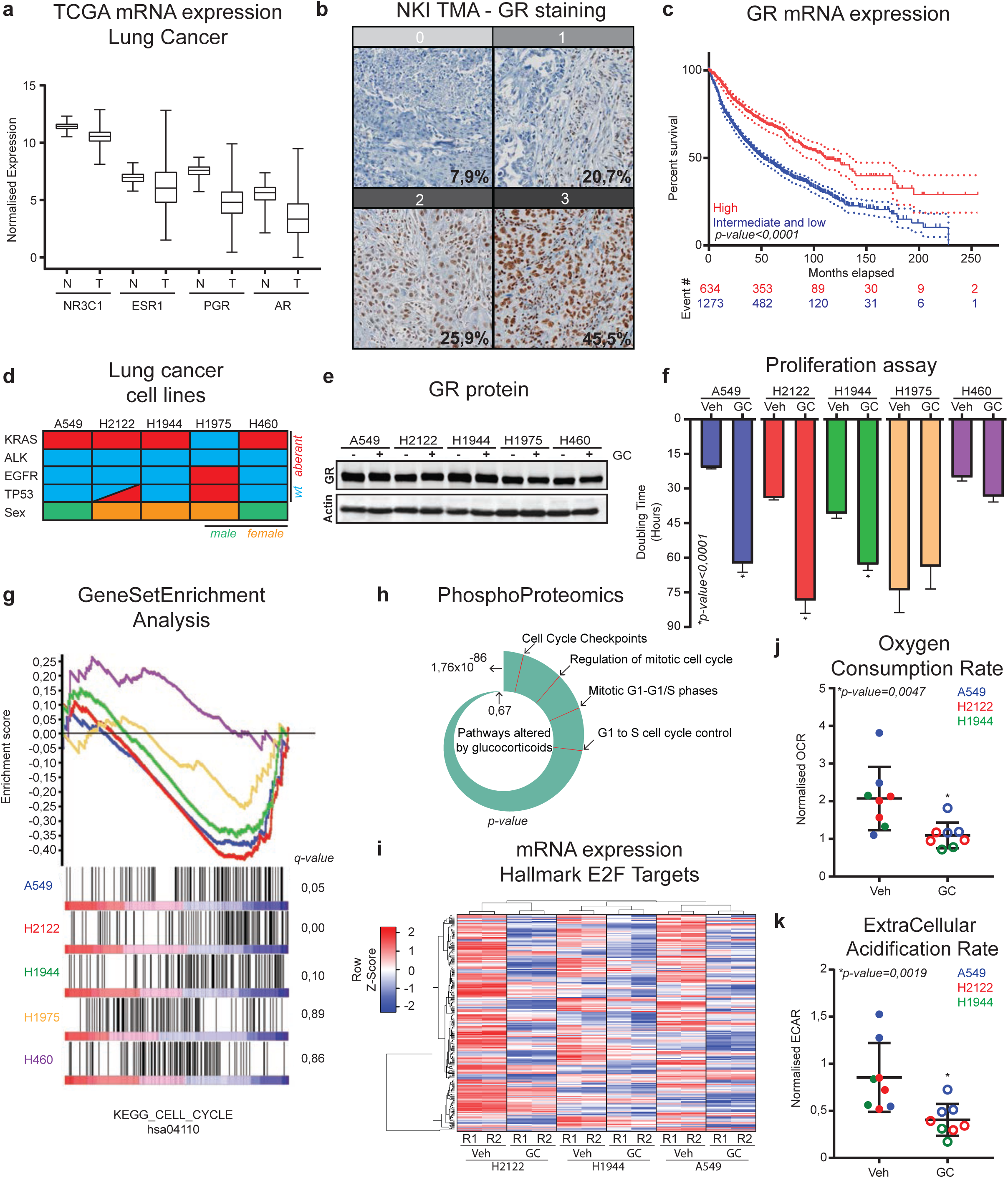
Glucocorticoid receptor induces quiescence in lung cancer models. **a,** Normalised mRNA expression levels of steroid hormone receptors across TCGA cohort in normal (N; 110 samples) and tumour (T; 1017 samples) tissue. **b,** Immunohistochemistry of GR in lung cancer. Representable immunohistochemistry images from low (0) to high (3) expression. **c,** Kaplan-Meijer curves based on GR mRNA expression, separated based on levels (high vs intermediate/low; 637 vs 1279 patients). **d,** Characteristics of cell lines used in the study. **e,** Representative western blot showing expression of GR, with actin as loading control. n=2. **f,** Doubling time (in hours) as calculated from real-time proliferation data of cells treated with GCs or vehicle. Bars depict mean ±SEM. n=4. **g,** GSEA for cell cycle gene set (hsa04110) based on two RNA-seq replicates. **h,** PhosphoPath pathway analsysis depicting pathways altered by GC treatment across the responders. Cell cycle related pathways are labelled in red. n=3 **i,** Heatmap showing expression of E2F target genes (M5925 geneset) in three responders, with or without GC treatment. n=2. **j,** Per DNA content normalised oxygen consumption rate (OCR) for three responders, without (Veh) or with glucocorticoids (GC). n≥2, per cell line. Bars depict mean ±SEM. **k,** Per DNA content normalised extracellular acidification rate (ECAR) for three responders, without (Veh) or with glucocorticoids (GC). n≥2, per cell line. Bars depict mean ±SEM.

The steroid hormone receptor expression profiles found in human tumours were entirely recapitulated in *in vitro* models of lung cancer (Fig. 1d; A549, H2122, H1944, H1975, and H460 cell lines), with GR (*NR3C1*) being the only receptor consistently expressed across the panel on mRNA level (Fig. S1b). GR expression was confirmed by western blot analysis (Fig. 1e), demonstrating comparable expression levels across five cell lines. More importantly, GC treatment of A549, H2122, and H1944 (hereinafter collectively referred to as “responders”) led to a drastic reduction of proliferation rate without signs of apoptosis, as observed by live-cell tracking experiments using SiR-DNA (Fig. 1f). Conversely, H1975 and H460 (hereinafter collectively referred to as “resistant”) cell lines were unaffected by GC therapy. In addition to the *in vitro* data, GC-induced growth-arrest persisted *in vivo*, as shown by 20-day mouse subcutaneous xenograft intervention study using H1944 cells (Fig. S1c).

In agreement with the observed growth-arrest upon GC treatment, strong devaluation of cell cycle genes (KEGG pathways; M7963) on whole transcriptome level was observed following activation of GR by GCs, exclusively in the responders (Fig. 1g). This was confirmed by PI-staining and flowcytometry experiments in which a reduction in S-phase and increased G0/G1-phase of the cell cycle occurred solely in the responders (Fig. S1d). In agreement with this, alterted phoshphorylation status of various cell cycle proteins (including Rb) (Fig. 1h and S1e) and marked downregulation of E2F-target genes following GR activation were observed (Fig. 1i). Upon GC treatment, the responders underwent distinct morphological changes, and stained positive for senescence associated X-gal (Fig. S1f). In addition, GC treatment lead to a significant decrease in oxygen consumption rate (Fig. 1j and S1g), and extracellular acidification rate (Fig. 1k). Together all of these features indicate responders underwent transition to a quiescent state upon GC treatment^13^. The same observations were found in both primary patient-derived and pre-established models of mesothelioma^14^; a cancer type derived from cells of mesodermal lineage (Fig. S1h-k).

### *CDKN1C* drives glucocorticoid-induced cell dormancy

In order to identify the downstream effector driving GC-induced cell cycle arrest, we performed RNA sequencing in the responders treated *in vitro* with vehicle or GCs for eight hours. GR was found to regulate approximately 500 genes across cell lines, with the transcriptional program partially shared (Fig. 2a). Focusing analysis on the upregulated genes (*log2fold≥2* and *padj≤0,01*) revealed 63 genes shared (Fig. 2b); with only one being a cell cycle regulator (*CDKN1C, encodes for* p57). In addition, this gene was also found upregulated in H2795 mesothelioma cell line which was growth-arrested by GCs, but not in two GC-resistant mesothelioma models (Fig. S2a). Supporting the observation that *CDKN1C* is regulated by GCs *in vivo*, a significant degree of Pearson correlation was found between classic GR-regulated genes (*PER1, NFKBIA, STOM, FKBP5,* and *DUSP1*) and *CDKN1C* across different human organ systems, whereas an absence of correlation with *ACTA1* mRNA expression was observed (Fig. S2b). In addition, analysis of human lung cancer specimens revealed that the *CDKN1C* gene was methylated and lowly expressed, in comparison to normal tissue (Fig. 2c).

**Figure 2.**
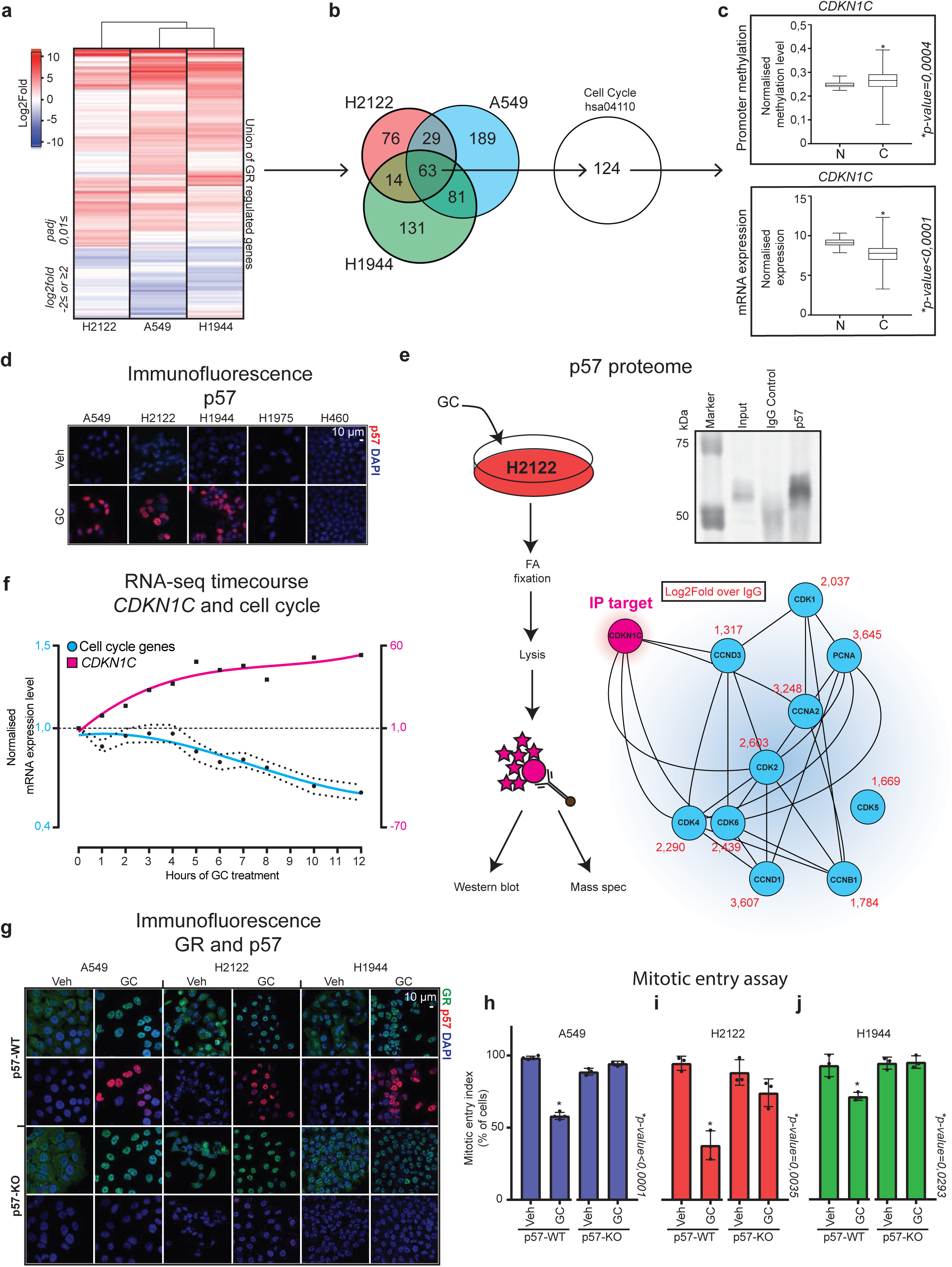
p57 is essential for GC-induced growth arrest. **a,** Complete lineage Euclidean clustering of GR regulated genes across three cell lines that are growth arrested by GCs. **b,** Intersect of genes differentially upregulated by GCs with a cell cycle geneset. **c,** Normalised *CDKN1C* promoter methylation (top; 75 vs 830 samples) and RNA expression (bottom; 110 vs 1017 samples) in normal (N) and tumour (T) samples from TCGA data. Bars depict min to max. **d,** Representative immunofluorescence images showing expression and localization of p57 (red), with DAPI as nuclear staining (blue). Scale bar, 10 μm. n=3. **e,** Scheme of IP experiment (left), and western blot for p57 (top right). Pathway analysis showing cell cycle genes identified by mass spectrometry analysis (bottom right). The values in red are log2fold differences in comparison to IgG control. Connecting lines indicate previously reported link. n=4. **f,** Normalised RNA expression level throughout the time-course experiment with glucocorticoids (GC) in A549 cells (ENCSR897XFT). Left axis depicts values for cell cycle genes, right axis depicts values for *CDKN1C*. n=3. **g,** Representative immunofluorescence images showing expression and localization of GR (green), p57 (red) in p57-WT and p57-KO cell lines. DAPI is used as nuclear staining (blue). Scale bar, 10 μm. n=3. **h-j,** Percentage of cells undergoing mitosis in a 60-hour experiment under vehicle (veh) and glucocorticoid (GC) treatment. n≥3. Bars depict mean ±SD.

Subsequently, levels of p57 were analysed by immunofluorescence and western blot. GC-dependent expression of p57 and its nuclear localization were found exclusively in the responders, whereas the expression was completely absent in the resistant cell lines (Fig. 2d and S2c). Furthermore, the protein interactome of p57 was profiled using mass spectrometry. These data confirm and expand a well-established protein interactome of p57, which is known to bind several cyclins and cyclin dependent kinases (CDKs)^15^, as well as PCNA^16^. In line with its function, a rapid and stable upregulation of *CDKN1C* mRNA levels by GC treatment preceded decreased expression of various cell cycle regulators (Fig. 2f), implying an existence of a transcriptional feedback mechanism.

To illustrate whether p57 is the sole driver of GR-induced growth-arrest, we performed CRISPR-Cas9 mediated disruption of *CDKN1C* in the responders. While GR function was unaffected, as seen through its ability to translocate to the nucleus following GC-treatment, p57 expression induction in a polyclonal population was almost entirely absent (Fig. 2g, *p57-KO*). To inspect whether GCs are still able to induce growth-arrest in p57-knockouts (p57-KO) live-cell imaging of SiR-DNA stained cells with and without GCs was performed, and the number of cells undergoing mitosis in the first 60 hours of treatment quantified. Disruption of the *CDKN1C* gene was enough to completely rescue the cells from GC-induced growth arrest (Fig. 2h-j). In addition, senescene associated X-gal staining was performed after treatment with vehicle (Veh) and GCs in H2122 p57-WT and p57-KO. Even though morphological change of the cells was observed upon GC treatment, the X-gal staining was entirely absent in p57 knockout cells (Fig. S2d), suggesting that the quiescence program was not initiated, which is in line with previously published work suggesting that p57 is essential for quiescence^17^.

### Glucocorticoids induce dependency on IGF-1R signalling through regulation of *FOXO1* and *IRS2*

As acquisition of cell dormancy is linked to reprogramming of signalling cascades^18^, we performed a drug screen (>2000 compounds) in the H1944 cell line to identify acquired pathway dependencies that arise when cells are pre-treated or co-treated with GCs (Fig. 3a, b).

**Figure 3.**
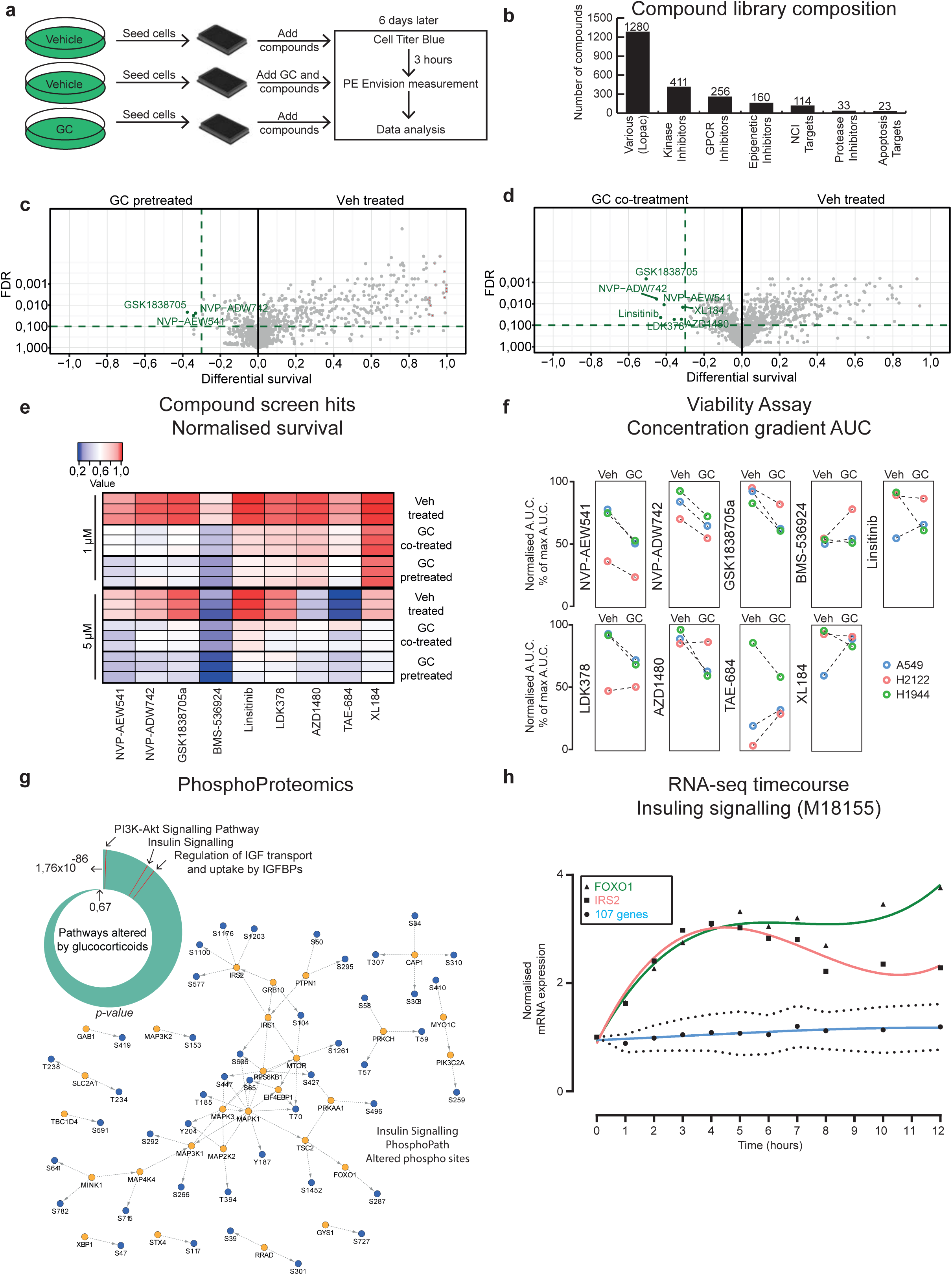
Drug screen identifies dependency on *IRS2-FOXO1* upon GC treatment. **a,** Set-up of the drug screen in H1944 cells. **b,** Categorisation of the compounds used in the screen. **c,** Volcano plot showing comparison of GC-pretreated arm and vehicle (Veh) treated arm. Compounds giving significantly differential survival in the GC-pretreated arm are depicted in green (*FDR≥0,1 and Differential survival≤0,35*). **d,** Volcano plot comparing GC co-treatment arm with vehicle control (Veh). Significantly differential survival in GC co-treatment arm depicted in green (*FDR≥0,1 and DF≤0,35*). **e,** Heatmap of normalised survival values for significant screen hits. Maximal survival is red (1), while blue represents minimal survival (0). **f,** Drug screen validation in A549, H1944, and H2122 cell lines. Area under the curve (AUC) values of the concentration gradient are normalised to maximal AUC. Two biological replicates were performed for all, except for H2122 treated with GSK1838705c where one biological replicate is shown. **g,** PhosphoPath pathway analysis identified insulin signalling related pathways enrichmed in GC treated cells (A549, H2122, and H1944). Differentially phosphorylated proteins within the “Insuling Signalling” pathway are depicted in orange and corresponding phospho-sites shown in blue. n=3. **h,** Normalised RNA expression levels of *FOXO1*, *IRS2*, and other genes of the insulin signalling geneset (M18155) throughout the time-course experiment with glucocorticoids (GC) in A549 cells (ENCSR897XFT). n=3.

Strikingly, treatment with GCs induced resistance to numerous drugs used in the screen (Fig. S3a). This further strengthens the conclusions made on a limited number of compounds by Herr *et. al*^6^, suggesting that widespread use of GCs in clinical management of cancer patients may reduce treatment efficacy. In contrast, GCs increased sensitivity to only a few inhibitors. More specifically, all identified compounds with a significant degree of synergism with GCs were IGF pathway inhibitors (*FDR<0,1* and *log2fold<0,35*) (Fig. 3c-e). Eight out of nine compounds were successfully validated in H1944 cells (Fig. 3f), and most of these were co-validated in A549 cells (Fig. 3f). As seen in the data available on The Cancer Dependency Map portal, the H2122 cell line is dependent on IGF1R signalling in absence of GCs (Fig. S3b), which explains why only slight synergism was found (Fig. 3f). In conjunction with this, phosphoproteomics-mass spectrometry experiments revealed that insulin signalling/IGF and the corresponding downstream pathways had altered phosphorylation status upon GC treatment (Fig. 3g).

Prior work has implicated GR in a cross-talk with mTOR in muscle tissue physiology^19^ and recent data suggest that insulin/IGF signalling is tightly bound to the circadian rhythm^20^. Based on this, we explored a GC-treatment time-course RNA-seq dataset, to reveal potential regulation of the insulin/IGF pathway by GCs. Out of all insulin/IGF signalling pathway genes, only *FOXO1* and *IRS2* were quickly and stably upregulated by GCs (Fig. 3h). In relation to GR regulation of *FOXO*1 and *IRS2*, we found positive Pearson correlation with mRNA expression of canonical GR target genes (Fig. S3c, d), indicating that these genes are regulated by GCs in normal physiology. Cumulatively, these data show that GCs induce IGF pathway dependency, which in turn reprograms cellular networks through diverse downstream mechanisms^21^.

### *Expression of CDKN1C*, *FOXO1*, and *IRS2* is directly regulated by GR through long-distance enhancer-promoter looping

To explore how GR regulates *CDKN1C, FOXO1,* and *IRS2* we have performed various techiques to explore the 3-dimensional chromatine organization in their respective vicinities.

We performed Circularized Chromosome Conformation Capture (4C)^23^ experiments to unbiasedly identify potential enhancer regions of the *CDKN1C* gene. In the responders, two potential distal enhancer regions (*a* and *b*) were identified which looped and contacted the *CDKN1C* promoter upon GC treatment (Fig. 4a). Induction of looping was entirely absent in the resistant cell lines (Fig. 4a). Both potential enhancer regions were found in a topologically associating domain (TAD) containing a large part of the *KCNQ1* and *CDKN1C* genes, flanked by CTCF sites as seen through Hi-C and ChIP-seq analysis, respectively (Fig. 4b). To show that GR is the essential factor for GC-induced expression of p57, we generated GR knockout H2122 cells (GR-KO). In H2122 GR-WT cells upon GC treatment, nuclear localization of GR and a concomitant expression of p57 was observed, while in the polyclonal GR-KO cell population no signal for both GR and p57 was detected (Fig. S4a). Therefore, to inspect the direct regulation of *CDKN1C* gene by GR, binding to the enhancer regions was investigated by ChIP sequencing (Fig. 4c). Chromatin binding of GR to both enhancer regions *a* and *b* was observed in the responders. Within the enhancer region *a* we observed a strong binding of GR to one site (Enhancer 1) containing two potential GR response elements, while for the enhancer region *b* we observed two distinct GR binding sites (Enhancer 2 and 3) located in close proximity to one another (Fig. 4c). Binding of GR to these sites was found in H1975, however it was absent in the H460 cell line. To address if the loss of binding was due to genomic deletions, we performed copy number analysis of shallow sequencing data. Although numerous different copy number alterations were found in the cell lines used, no copy number alterations (gains or losses) were detected in chromosome 11 (Fig. S4b). Interestingly, Enhancer 1 was marked by H3K27Ac, an active enhancer chromatin mark (Fig. S4c), while only a weak signal was seen for Enhancers 2 and 3 (Fig. S4d). Complementary to the 4C data (Fig. 4a), cohesin (SMC3/Rad21) recruitment was observed exclusively to Enhancer 1 (Fig. S4c and S4d). The strong H3K27Ac mark and cohesin recruitment, both suggest that Enhancer 1 is the main regulatory element through which GR regulates *CDKN1C* gene expression, while Enhancers 2 and 3 may serve as auxiliary enhancers.

**Figure 4.**
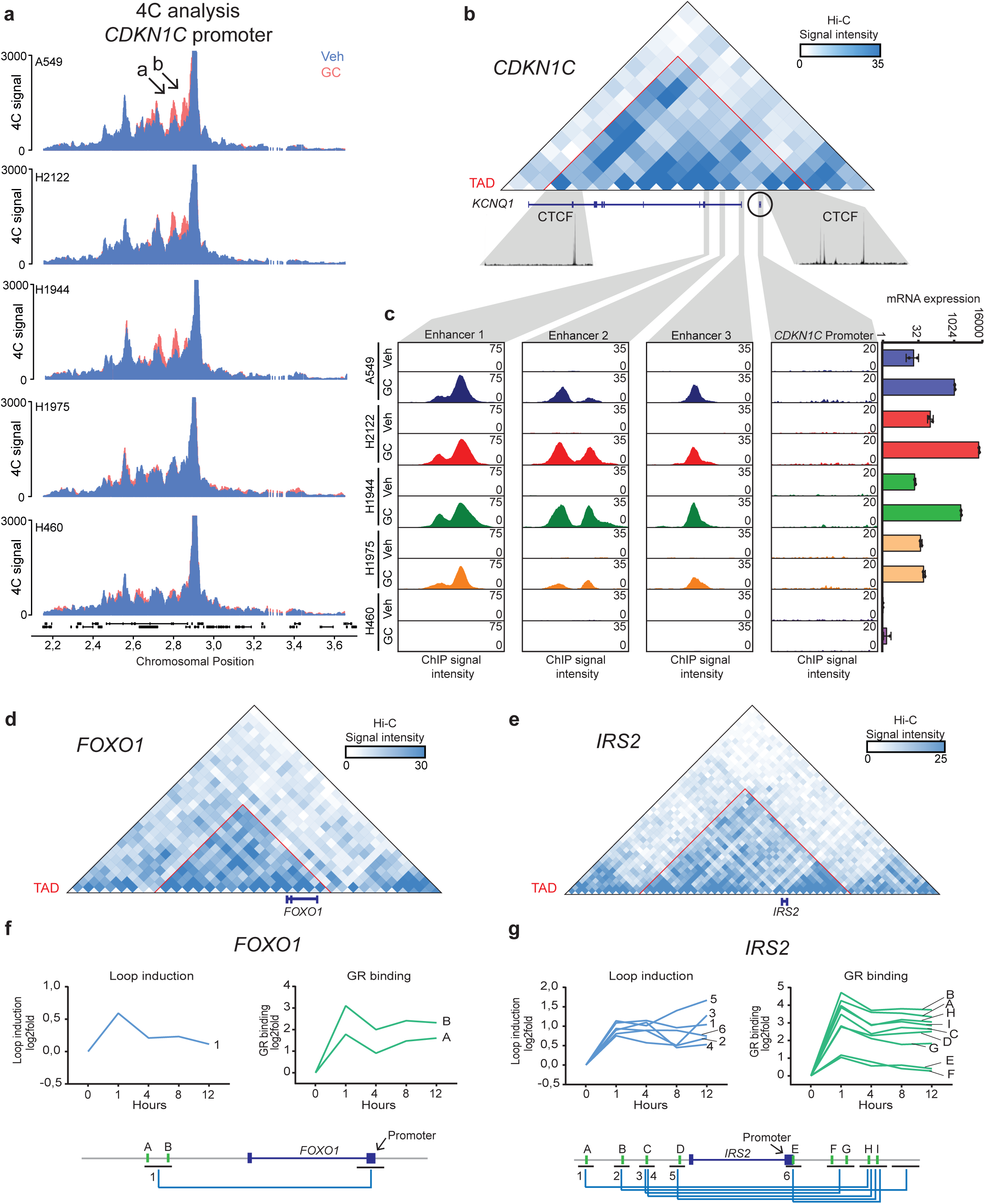
Expression of p57 is regulated through genetic looping induced by GR binding to a distal enhancer. **a,** 4C signal across the region surrounding *CDKN1C* gene on chromosome 11, under Vehicle (Veh; blue) and glucocorticoid (GC; red) treatment. Average signal of 2 biological replicates shown. **b,** Hi-C contact map of A549 cell line at resolution of 40 kb of the region surrounding *CDKN1C* gene (chr11:2421230-3141230). CTCF ChIP data (U01HG00790) are represented for TAD anchor sites. **c,** GR ChIP-seq data showing peaks at enhancer 1/2/3 (chr11:2799339-2801363; chr11:2846111-2848121; chr11:2880971-2882981, respectively), and *CDKN1C* promoter (chr11:2900697-2910745) for all the cell lines, not-treated (Veh) or treated (GC). Right: mRNA expression of *CDKN1C*. n=2. **d,** Hi-C contact map of A549 cell line at resolution of 40 kb of the region surrounding *FOXO1* gene (chr13:39760000-41120000). **e,** Hi-C contact map of A549 cell line at resolution of 40 kb of the region surrounding *IRS2* gene (chr13:108480000-110680000). **f-g,** Loop induction calculated from Hi-C time-course data for every loop detected in the vicinity of the locus of interest in A549 cell line (top left). Differential GR binding within loop anchors (compared to 0 hrs) across a 12hr time-course ChIP experiment in A549 cell line (top right). Schematic representation of the respective locus (bottom).

To inspect the direct regulation of *FOXO1* and *IRS2* we made use of GR ChIP-seq and Hi-C timecourse data from A549 cells. GR chromatin binding to several enhancer sites in the corresponding TAD (Fig. 4d, e) containing the *FOXO1* or the *IRS2* gene was observed (Fig. 4f, g). In addition to binding of GR to these enhancers, induction of a single enhancer-promoter loop containing two GR binding sites in the loop anchor for the *FOXO1* gene was observed (Fig. 4f). As for *IRS2*, a complex web of six enhancer-promoter loops was detected, containing nine GR binding sites (Fig. 4g). Collectively, regulation of *CDKN1C*, *FOXO1*, and *IRS2* resembles the previously proposed model of gene regulation by the oestrogen receptor^24^, however the enhancer distance by far exceeds previously reported general modes of regulation.

### SWI/SNF complex forms an integral part of a proficient GR transcriptional machinery

Resistance to GCs can arise through alterations in any of the multiple nodes in the GR signalling axis^25^. We have therefore examined what drives GC resistance in the cell culture models used in this study. For all five cell lines used, GR was able to effectively translocate to the nucleus in response to GCs (Fig. 5a) and bind the chromatin (Fig. 5b), both in responders and resistant cell lines. Further, we performed RNA-seq experiments to test whether GR is able to modulate gene expression across the five cell lines used. We observed a high degree of gene expression modulation upon GC activation exclusively in the responders, while in the resistant cells only few genes were significantly affected (Fig. 5c).

**Figure 5.**
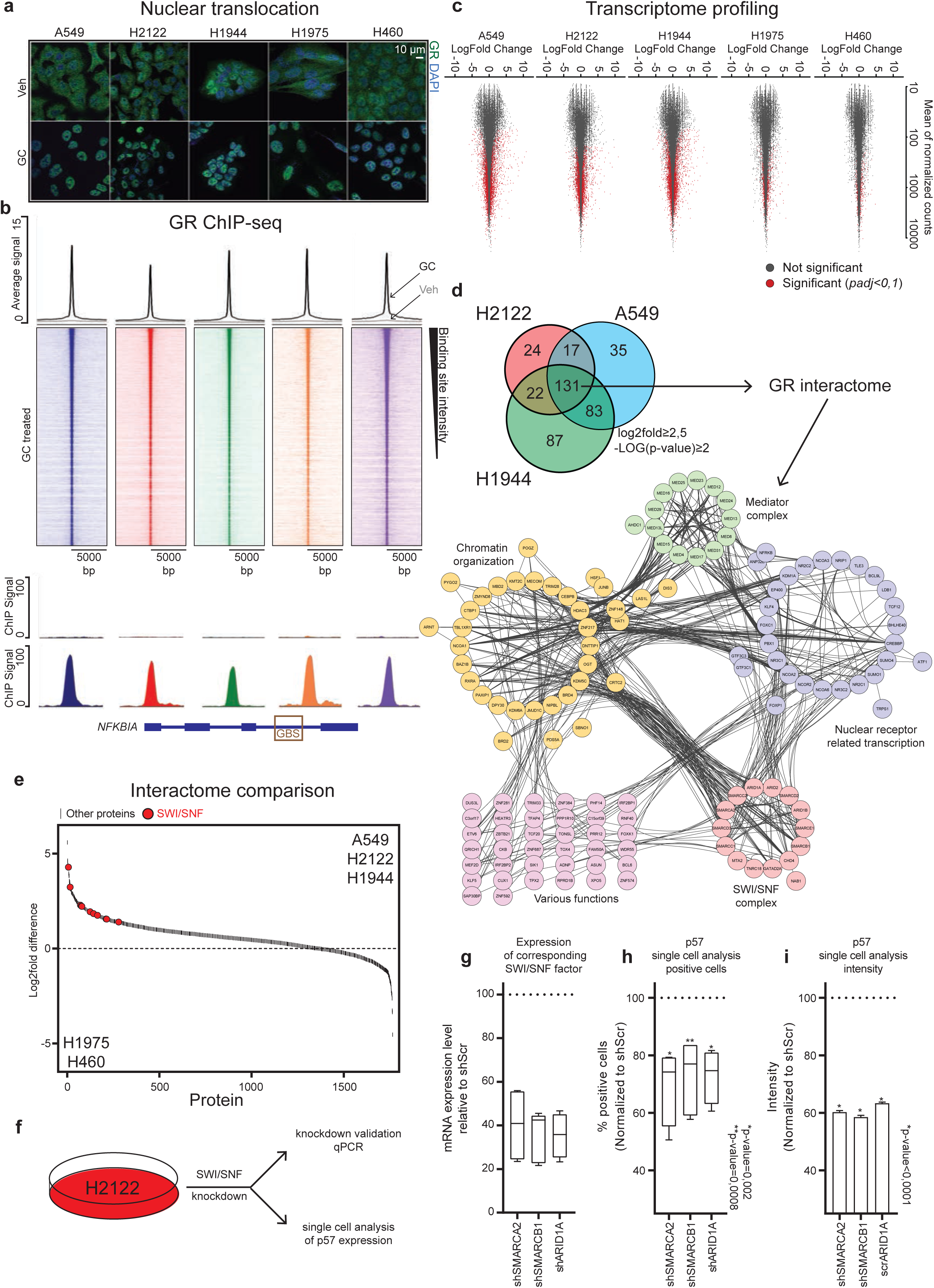
SWI/SNF complex is a part of a proficient GR transcriptional machinery. **a,** Representative immunofluorescence images showing expression and localization of GR (green), treated with glucocorticoids (GC) or control (Veh), using DAPI as nuclear staining (blue). Scale bar, 10 μm. n=3. **b,** Average signal of GR ChIP sequencing experiments across all sites called over input control (top) and coverage plot visualization of all sites (middle). Signal at classical GR regulated gene *NFKBIA* across all cell lines treated with glucocorticoids (GC) or control (Veh) (bottom). **c,** MA-plot depicting differential gene expression induced by GC treatment in RNA-seq experiment. Genes significantly (*padj≤0,1*) up- or down-regulated by GCs are depicted in red. n=2. **d,** Intersect of all GR interacting proteins found by mass spectrometry (*log2fold≥2,5* and *-LOG(p-value)≥2*) between three cell lines growth arrested by GCs (top). String pathway analysis of all common interactors. Lines represent previously reported links. n=3. **e,** Waterfall plot depicting differentially enriched interactors between three responding and two resistant cell lines after GC treatment. SWI/SNF complex members are depicted in red. n=3. **f,** Scheme showing the SWI/SNF knockdown experiments. For each gene *≥2* different shRNAs were used. Each experiment was performed in 2 biological replicates. **g,** Normalised mRNA expression level relative to shScr control for *SMARCA2*, *SMARCB1*, and *ARID1A*. Bars depict min to max. **h,** Percentage of cells stained positive for p57 normalised to shScramble control. Bars depict min to max. **i,** Intensity level of p57 staining shown in percentage of shScramble control. Bars depict mean ±SEM.

Based on the results described above we hypothesized that differences in the GR protein complex may underpin the absence of response to GC treatment. Therefore, we performed rapid immunoprecipitation mass spectrometry of endogenous proteins (RIME)^26^ to compare GR transcriptional complex composition in the responders versus resistant cells. Analysis of enriched interactors in three responders enabled us to annotate the common GR transcription complex members (Fig. 5d). The common GR interactome can be divided into four distinct categories based on pathway enrichment analysis: nuclear receptor related transcription, chromatin organization, mediator complex, and SWI/SNF complex. Interestingly, on the phosphoproteomics level we detected numerous modifications in the chromatin organization pathway (including SWI/SNF) (Fig. S5a), suggesting a cross-regulatory network with GR. Statistical comparison of RIME data of responder and resistant cell lines revealed differences in the GR complex (Fig. 5e). Among the most-depleted GR interaction partners in the resistant cells were the members of the chromatin remodelling complex (SWI/SNF) (Fig. 5e). As distinct components of the SWI/SNF complex were previously related to GR signaling^27^, with low levels being associated with GC resistance in acute lymphoblastic leukemia^28^, we explored whether SWI/SNF loss would affect GR signalling. For this, we used the H2122 cell line and p57 protein expression as a proxy for GR activity. Using at least two shRNAs per target (*SMARCA2*, *SMARCB1*, and *ARID1A*) we perturbed the expression of specific SWI/SNF components. The degree of knockdown was checked using qPCR, confirming >50% knockdown efficacy for all three SWI/SNF components (Fig. 5g). Quantifying the p57 signal in over 10.000 individual cells revealed a significant decrease in both the number of p57 positive cells (Fig. 5h) and the staining intensity (Fig. 5i) upon loss of SWI/SNF. With this, we have shown that SWI/SNF forms an essential part of the GR transcriptional complex, necessary for regulation of p57, which is required to drive cells into quiescence.

### *CDKN1C*, *FOXO1*, and *IRS2* form a tumour suppressor axis in human cancer which can be inactivated through alterations in SWI/SNF or TP53

*CDKN1C*, *FOXO1*, and *IRS2* have previously been implicated in differentiation and maintenance of cellular identity^29–31^. To inspect whether the *CDKN1C*/*FOXO1*/*IRS2* axis is relevant for tumour development and/or progression, we explored mRNA sequencing data across 32 human cancer types (9874 samples). A significant decrease in expression of all three GR-regulated genes was evident between normal and cancer samples (Fig. 6a). Unsupervised hierarchical clustering (Fig. 6Sa) of primary cancer samples based on mRNA expression revealed distinct clusters, among which a cluster with preserved expression of *CDKN1C*, *FOXO1*, and *IRS2* (normal-like), and a cluster with a loss of expression of all three (triple-loss) (Fig. 6b). Inspection of expression levels of tumour aggressiveness markers (*MKI67*, *FEN1*, *AURKA*, and *FOXM1*) between these two clusters revealed significant differences, pointing towards an aggressive molecular phenotype in triple-loss tumours across all cancer types (Fig. 6c). The same analysis was applied to datasets containg only lung or breast cancer samples and yielded identical observations (Fig. S6b-e).

**Figure 6.**
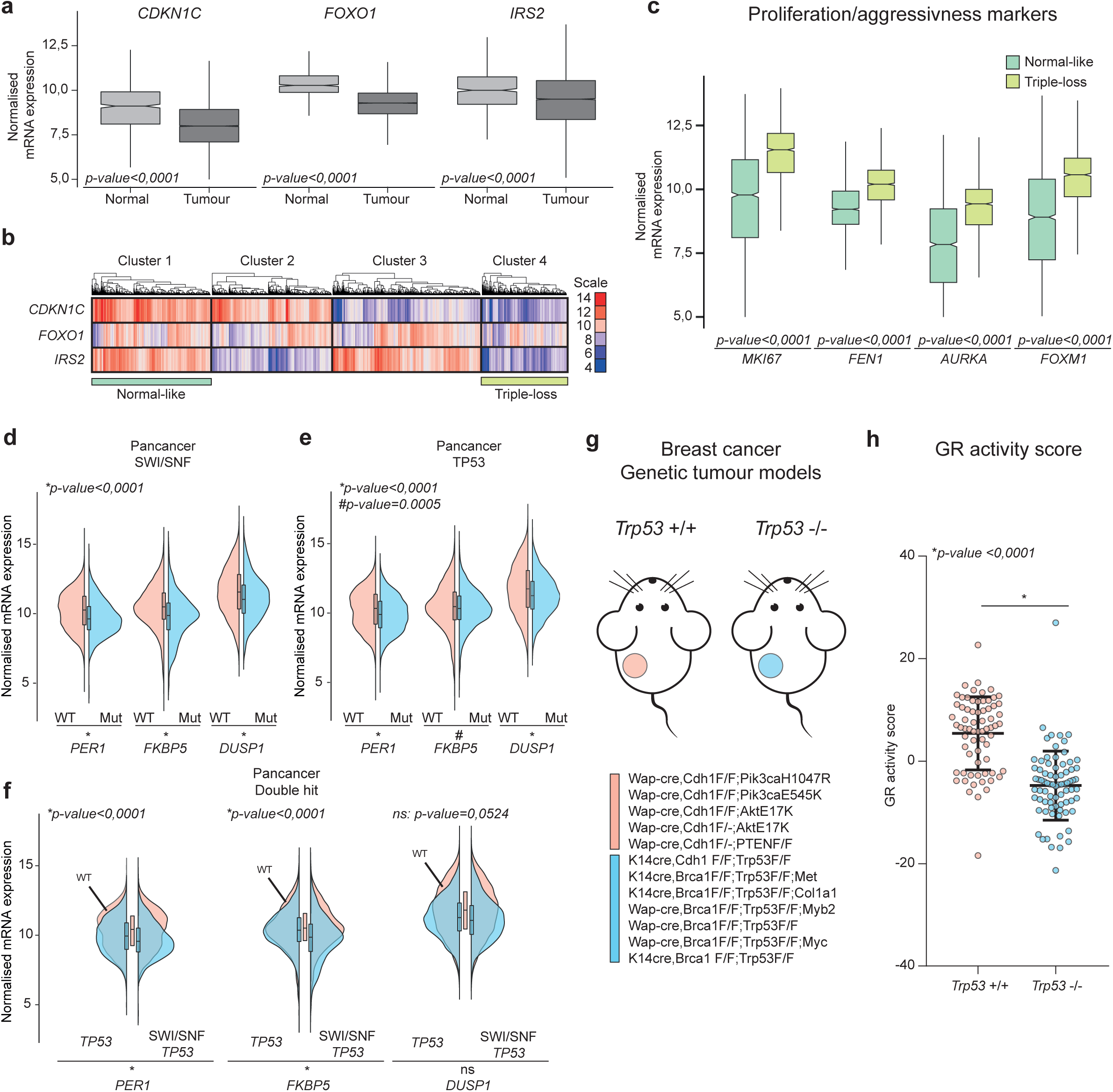
*CDKN1C, FOXO1,* and *IRS2* in human cancers and GR inactivation by SWI/SNF and *TP53* alterations *in vivo*. **a,** Bar plots showing normalised mRNA expression of *CDKN1C*, *FOXO1*, and *IRS2* across normal (light grey) and tumour (dark grey) specimens (top). **b,** Unsupervised clustering (k_means_=4) of 9874 cancer samples. **c,** Normalised mRNA expression of proliferation/aggressiveness markers (*MKI67*, *FEN1*, *AURKA*, and *FOXM1*) between ‘normal-like’ and ‘triple-loss’ cluster. **d-f,** Violin plots of mRNA expression levels of three classic GR regulated genes in samples harbouring mutations or unaffected loci (WT) across 32 tumour types. **g,** Schematic representation and list of genetic breast cancer mouse models used. **h,** GR activity score for *Trp53*-WT (pink) or *Trp53*-KO (blue) mouse mammary cancers.

Inactivation of genes involved in tumour suppression and/or cell dormancy can occur through various alterations, on both the genetic and epigenetic level^32^. Genetic alterations of the *NR3C1* gene (copy number changes and mutations) are rare in cancer (Fig. S6f). In relation to our previous experiments (Fig. 5h, i) we hypothesized that SWI/SNF (*SMARCA2*, *SMARCB1*, and *ARID1A*) mutations may deactivate the GR axis. In addition, we hypothesise that *TP53* can also contribute to inactivation of GR signalling, as it is a well-established crucial regulator of quiescence^33,34^ and crosstalk with GR was previously proposed^35^.

We therefore explored whether mutations in both SWI/SNF and *TP53* contribute to inactivation of the GR signalling axis. This was achieved by inspecting expression levels of three classic GR regulated genes (*PER1*, *FKBP5*, and *DUSP1*) across 32 different tumour types. Mutations in SWI/SNF components were associated with significant downregulation of the three target genes tested (Fig. 6d), providing *in vivo* confirmation of our experimental findings (Fig. 5h, i). Similarly, *TP53* mutations negatively correlated with GR target gene expression (Fig. 6e). The data also suggest that SWI/SNF and *TP53* act via different mechanisms on GR signalling, as co-occurrence of SWI/SNF and *TP53* mutations illustrated an additive effect on GR target gene downregulation across tumour types (Fig. 6f). Importantly, using twelve different genetically engineered mouse models of breast cancer (Fig. 6g), we show that *Trp53* loss in mammary tumours leads to a severe dysfunction of the GR signalling axis (Fig. 6h), as seen through significant decrease in the GR activity score, which was computed based on genes that are direct targets of GR in a non-malignant epithelial breast cell line^36^.

## Discussion

Maintenance of cellular integrity and differentiation programs is achieved through various protein-protein and protein-DNA interactions. A multitude of experimental and empirical findings point to the existence of a redundant network of molecules supporting cellular homeostasis, in turn suppressing loss of lineage features and abnormal growth. With advancements in sequencing technologies and increasing number of patient samples processed, various proteins joined the ranks of the tumour suppressive network with *RB1* and *TP53*^37^.

Steroid hormone receptors have been in the spotlight of cancer research for decades, yet their role in normal tissue homeostasis is still not entirely understood. For example, the androgen receptor is considered an oncogene, driving growth and resistance of prostate cancer^38^. However, studies suggest that in terms of prostate development AR operates as a tumour suppressor^39^. Using a comprehensive multidisciplinary approach, we have shown that GR through induction of chromatin looping, regulates cell cycle arrest (through p57) and insulin/IGF signalling pathway (via *FOXO1/IRS2*). We hypothesize that this feature of GR to induce cellular dormancy (quiescence) may be reminiscent of its signalling in normal tissue. In conjunction with this hypothesis, activation of GR has been previously linked to cell differentiation and lineage selection^40,41^. As tumours are exposed to GCs from the circulation, this dormant state might be a feature of various early-stage human tumours. Quiescense has been shown to be under the circadian control in various tissues and stem cells^42,43^. In relation to that, the well-known day-night rhythmic behaviour of GC levels may also suggest that quiescence in cancer, and therefore sensitivity to chemotherapy, is subjected to circadian rhythm through GR activity. In fact, morning-evening differences in chemotherapy response in leukaemia patients have been observed over 30 years ago^44^.

In order to mediate quiescence GR needs to form a transcriptionally proficient complex. We have shown that this complex consists of various proteins involved in transcriptional regulation (transcription factors and mediator complex), as well as factors involved in genome organization. The recruitment, physical association or proximity of GR to the genome organization machinery (including the SWI/SNF complex) as reported in this manuscript, is striking. These interactions would enable modulation of gene expression through reorganization of the chromatin in three-dimensional space and further support the notion that GR could act as a pioneering factor as proposed previously by others^45^. Additionally, our data suggest that interaction of GR with the chromatin organization machinery extends beyond recruitment and physical proximity, as GC treatment led to changes in their posttranslational modifications (phosphorylation). Altered phosphorylation status of SWI/SNF family^46^ and chromatin modifiers^47,48^ has been linked to modulation of their activity. Therefore, changes in posttranslational modifications that occur by GC treatment might contribute to the pioneering potential of the GR.

Factors involved in cell dormancy (such as *TP53* and SWI/SNF components) may be inactivated through genomic alterations; however, the GR gene locus is not disrupted in cancer. Strikingly, we have shown that SWI/SNF loss, as well as *TP53* deficiency, have deleterious effects on the GR axis. These data suggest that the GR axis operates as an integral part of a larger signalling network^32^, probably downstream of *TP53* and SWI/SNF, to co-ordinate cell cycle progression and lineage selection. Importance of GR in this network is further supported by an *in vivo* the GR haploinsufficiency model, in which loss of a single copy of GR was enough to induce various malignancies across different organs^4^.

The mechanistic insight into the GR-induced quiescence provides a possible explanation to why adrenocortical carcinoma (ACC) is the only tumour type in which IGF1R inhibitor Linsitinib had a significant effect^49^. High levels of intratumoral cortisol can be found in ACC, in addition, loss of *CDKN1C* expression/function has been linked to development or progression in ACC^50^. One could hypothesize that in absence of p57 expression, tumours with high GR activity, due to high local cortisol levels, would be dependent on IGF signalling route. In turn, this would enable eradication of ACC cells through inhibition of this axis. Future research should address whether this approach could be used for treatment of other solid tumours with high GR activity and loss of p57.

Our findings can be extrapolated on a wider scale suggesting that GR, with its target genes, forms an axis protecting various tissue types of cancer development or in terms of early primary cancers keeping the cells in a dormant state. This axis would tie, together with *TP53* and SWI/SNF, into an expanding network of proteins operating towards maintenance of cellular integrity and differentiation programs throughout human physiological systems.

## Acknowledgment

This work was funded by the Netherlands Organization for Scientific Research NWO VIDI grant 91716401, an Alpe d’Huzes/KWF Bas Mulder Award, and Oncode Institute. Joana Silva was supported by an EMBO long-term fellowship (ALTF 210-2018). We would like to acknowledge the NKI-AVL Core Facility Molecular Pathology & Biobanking (CFMPB) for lab support, the NKI Genomics Core Facility for Illumina sequencing and bioinformatics support, and the NKI mouse intervention unit for performing xenograft experiments. We thank Marlous Hoogstraat for the copy number analysis pipeline; Laurel M. Schunselaar for help with lung cancer pathological samples; Suzan Stelloo and Simon Linder for frequent, and helpful discussions.

## Contributions

S.P. W.Z. conceived the project outline and coordinated the project. S.P. designed and implemented the project, W.Z. was responsible for project funding and supervision, S.P. K.S. I.M.P. performed RNA and ChIP sequencing experiments, S.P. K.S. A.H. performed flowcytometry experiments, A.H. generated all the data on mesothelioma cell lines, S.P. K.S. S.Y. generated and performed experiments on knockout/knockdown models, S.P. M.B. L.H. M.A. performed mass spectrometry experiments and analysis, A.G.M performed live-cell imaging experiments and analysis, S.P. K.S. H.T. E.dW. performed 4C experiments and analysis, A.B. T.R. analysed GR-ChIP and Hi-C time-course data, S.P. J.S. W.F. performed and analysed seahorse experiments, S.P. K.S. B.M. C.L. R.B. designed, executed, and analysed the drug screen, B.G. provided mRNA expression and associated clinical data of human lung cancer tissue, K.M. performed scoring of IHC tissue microarray, M.D.W. J.J. K.E.dV. generated breast cancer mouse models and performed the experiments, S.P. B.vdB. performed image analysis, S.P. W.Z. designed, supervised, and analysed mouse intervention experiments, S.P. performed computational analysis of majority of data, S.P. K.S. A.G.M. R.M. W.Z. discussed and interpreted the data, S.P. wrote the manuscript, with input from all authors.

## Materials and Methods

### Cell lines

A549, H2122, H1944, H1975, and H460 cells were obtained from Rene Bernard’s lab (Netherlands Cancer Institute, Netherlands). HEK293T cells were obtained from American Type Culture Collection (ATCC). The human mesothelioma cell lines M28 and VAMT were provided by Prof. Broaddus (University of California, USA), while the NCI-H2795 (H2795) was obtained from Prof. McDermott (Sanger Institute, UK). The primary human mesothelioma cell lines PV130913 and PV041214 were generated by Dr. Schunselaar (Netherlands Cancer Institute, Netherlands).

A549 and H2795 cell lines were maintained in DMEM/F12 (1:1) (1X) + Glutamax (Life Technologies), while HEK293T, M28, PV130913 and PV041214 were cultured in DMEM, both media supplemented with 10% foetal calf serum (FCS). The other cell lines were cultured in RPMI-1640 (phenol-red + glutamine) (Life Technologies) medium supplemented with 10% FCS. All the cell lines were authenticated and tested negative for mycoplasma contamination.

For hormone deprivation, cells were cultured in medium containing 5% charcoal-treated FCS for three days, subsequently treated with 2,75 µM hydrocortisone (Sigma) or vehicle, and harvested at the indicated time point.

### Generation of CRISPR cell lines

Guide RNAs targeting human *CDKN1C* (CTGGTCCTCGGCGTTCAGCT), *NR3C1* (GTGAGTTGTGGTAACGTTGC), and a non-targeting control guide (GTATTACTGATATTGGTGGG) were individually cloned into the lentiCRISPR v2 plasmid^1^. CRISPR vectors were co-expressed with 3^rd^ generation viral vectors in HEK293T cells using PEI. After lentivirus production, the medium was harvested and transferred to the designated cell lines. Two days post infection cells were put on puromycin selection for three weeks.

### shRNA experiments

shRNA knock-down experiments were carried out by infection of H2122 cells with pLKO.1-puro containing a non-targeting, SMARCA2-, SMARCB1-, or ARID1A-specific hairpin obtained from The RNAi Consortium (TRC) library (https://www.broadinstitute.org/rnai-consortium/rnai-consortium-shrna-library).

### Xenografts

The H1944 cells were trypsinised and resuspended in PBS at a density of 6 million cells/50 µl and mixed with an equal volume of BME (#3533-005-02, Sigma-Aldrich) NOD-scid gamma (NSG) mice (±7 weeks old) were anesthetised before injection of 6 million cells subcutaneously into one of the flanks. Once the tumour size reached between 250 and 300 mm^3^ the mice were treated with 75 mg/kg Dexamethasone (D2915-100MG, Sigma-Aldrich; dissolved in water) or vehicle by I.P. injections on a daily basis. Tumour volume was monitored by calliper measurements every day. The NKI Animal Experiments Committee approved all *in vivo* experiments (project number 9139).

### RNA sequencing

Cells were serum starved for three days before they were treated with hydrocortisone (2,75 µM) for 8 hours. Total RNA was isolated using the RNeasy Mini Kit (Qiagen, Germany) according to the manufacturer’s instructions. Quality and quantity of the total RNA was assessed by the 2100 Bioanalyzer using a Nano chip (Agilent, USA). Total RNA samples having an RNA integrity number (RIN) above 8 were subjected to library generation.

Strand-specific libraries were generated using the TruSeq Stranded mRNA sample preparation kit (Illumina, USA; RS-122-2101/2) according to the manufacturer’s instructions (Illumina, Part #15031047 Rev. E). 3′ adenylated and adapter ligated cDNA fragments were subject to 12 cycles of PCR. The libraries were analysed on a 2100 Bioanalyzer using a 7500 chip (Agilent, Santa Clara, CA), diluted and pooled equimolarly into a multiplexed, 10 nM sequencing pools and stored at −20 °C. Strand-specific cDNA libraries were sequenced with 100 base paired-end reads on a HiSeq2500 using V4 chemistry (Illumina). RNA sequencing data was mapped to exons using Tophat (v.2.1). Read counting, normalization, and differential gene expression was performed using R package DESeq2^2^.

### ChIP sequencing

Chromatin immunoprecipitations were performed as previously described^3^. Nuclear lysates were incubated with 7,5 μl of NR3C1 antibody (D6H2L, Cell Signalling Technology) pre-bound to 50 μl protein A beads per samples. Immunoprecipitated DNA was processed for library preparation (0801-0303, KAPA biosystems kit). Samples were sequenced using an Illumina Hiseq2500 genome analyser (65bp reads, single end), and aligned to the Human Reference Genome (hg19, February 2009). Reads were filtered based on MAPQ quality (quality ≥ 20) and duplicate reads were removed. Peak calling over input control was performed using MACS2 peak caller. MACS2 was run with the default parameters. Genome browser snapshots, heatmaps and density plots were generated using EaSeq (http://easeq.net)^4^.

### Western Blot

Cells were lysed in 2xLaemmli buffer (120 mMTris, 20% glycerol, 4% SDS). Total protein content was quantified by BCA assay (23227, Thermo Fisher Scientific). Cell lysates containing equal amounts of protein were analysed by SDS-PAGE, after protein transfer, nitrocellulose membranes were incubated with antibodies against GR (12041, Cell Signalling Technology,1:1000), p57 (sc-56341, Santa Cruz Biotechnology, 1:500), or Actin (MAB1501R, Merck,1:10.000).

### Seahorse

Cellular respiration was measured using a Seahorse XF24 Bioanalyzer (Seahorse Biosciences). A549, H2122, and H1944 cells were all seeded at 75,000 cells per well to a XFe24 cell culture microplates (102340-100, Seahorse Biosciences) and cultured overnight before the analysis. The analysis was performed according to the manufacturer’s instructions in DMEM (D5030, Sigma-Aldrich) supplemented with 10 mM D-glucose and 4 mM L-glutamine for the oxygen consumption rate (OCR) experiments. For OCR measurements the following reagents were added: oligomycin (1 μM), FCCP (0,4 μM), and rotenone (1 μM) and antimycin A (1 μM). Results were normalized to DNA content using nanodrop quantification (Thermo Fisher Scientific).

### Senescence associated X-gal assay

Cytochemical staining for senescence associated β galactosidase was performed as described before^5^. Cells were serum starved for three days and subsequently fresh medium with or without hydrocortisone (2,75 µM) was added for two days. After incubation, cells were washed twice with PBS and fixed with 3,7% formaldehyde for five minutes. Following that, cells were washed with PBS before they were incubated with X gal staining solution (1 mg/mL X-gal, 40 mM citric acid/sodium phosphate buffer, 5 mM potassium ferricyanide, 5 mM potassium ferrocyanide, 2 mM MgCl2, 150 mM NaCl) overnight at 37°C. X gal is an artificial substrate of the β galactosidase enzyme and is used to detect senescence associated β galactosidase. The next day, cells were washed with PBS and imaged with a Zeiss Axiovert S100 inverted microscope (Zeiss, Germany).

### Cell cycle analysis with flow cytometry

Cells were serum starved for at least three days and subsequently, either left untreated (FCS) or treated with hydrocortisone (2,75 µM) for 2 days. Subsequently, cells were harvested and centrifuged for 4 minutes at 1000 rpm at 4°C. The pellet was re-suspended in cold PBS and cells were fixed in cold 80% EtOH overnight at 20°C. The next day, cells were centrifuged for five minutes at 1500 rpm at 10°C and incubated with 500 μL of PI staining mix (50 μg/mL PI, 100 μg/mL RNAse A, 0,5% Triton™ X 100) for 40 minutes at 37°C. Following that, PBS was added, and samples were centrifuged for five minutes at 1500 rpm at 4°C. Pellets were re-suspended in PBS and stored at 4°C till flow cytometric analyses. Flow cytometric measurements were performed on LSRFortessa SORP 2 (BD BioSciences) and cell cycle distribution analysed with FlowJo Software (FlowJo LLC).

### Annexin V/Propidium Iodide apoptosis assay for flow cytometry

Annexin V/Propidium Iodide apoptosis assays were performed as described before^6^. Cells were serum starved for at least three days. Following that, cells were either left untreated (FCS) or treated with HC at a concentration of 2,75 µM for 6 days. As a positive control, cells were treated for approximately 30 hours with 50 µM cisplatin to induce apoptosis. Cells were harvested and centrifuged for 10 minutes at 335 rcf at 4°C. The pellet was washed twice, once with cold PBS and a second time with Annexin V binding buffer (10 mML HEPES, 140 mM NaCl, 2.5 mM CaCl2, pH 7.4). After centrifugation for 10 minutes at 335 rcf at 4°C, cells were resuspended in Annexin V binding buffer and Annexin V (Thermo Fisher Scientific, USA) was added according to the manufacturer’s recommendations. Samples were incubated for 15 minutes at room temperature in the dark before propidium iodide (PI) was added at a concentration of 2 µg/mL. Following an additional incubation for 15 minutes at RT, cells were washed with Annexin V binding buffer and centrifuged for 10 minutes at 335 rcf at 4°C. Cells were resuspended in Annexin V binding buffer with 1% formaldehyde and fixed for 10 minutes on ice or overnight at 4°C. Subsequently, cells were washed twice with cold PBS and centrifuged for 8 minutes at 425 rcf at 4°C. The pellets were resuspended in Annexin V binding buffer and RNase A (50 μg/mL; Thermo Fisher Scientific, USA) was added. Samples were incubated for 15 minutes at 37°C. and washed once more with cold PBS. Afterwards, cells were centrifuged for 8 minutes at 425 rcf at 4°C, re suspended in PBS and stored at 4°C till flow cytometric analyses. Flow cytometric measurements were performed on an Attune NxT Flow Cytometer (Thermo Fisher Scientific, USA) and cell populations analysed with FlowJo Software (FlowJo LLC, USA).

### RNA isolation and mRNA expression

Total RNA was isolated using TRIzol Reagent (Thermo Fisher Scientific, USA), and cDNA was synthesized from 2 μg RNA using the SuperScript™ III Reverse Transcriptase system (Thermo Fisher Scientific, USA) with random hexamer primers according to the instructions provided by manufacturers. Quantitative PCR (qPCR) was performed using the SensiMix™ SYBR Kit (Bioline, UK) according to the manufacturer’s instructions on a QuantStudio™ 6 Flex System (Thermo Fisher Scientific, USA). Primer sequences for mRNA expression analysis are listed in Supplementary table 1.

### Immunohistochemistry

Immunocytochemistry of formalin fixed paraffin embedded (FFPE) cell lines and immunohistochemistry of the FFPE tumour samples were performed on a BenchMark Ultra autostainer (ERα, PR, AR) or Discovery Ultra autostainer (GR). Paraffin sections were cut at 3 µm, heated for 28 minutes at 75°C and deparaffinized in the instrument with EZ prep solution (Ventana Medical Systems). Heat-induced antigen retrieval was carried out using Cell Conditioning 1 (CC1; Ventana Medical Systems, USA) for 32 minutes (AR), 36 minutes (ERα, PR) or 64 minutes (GR) at 95°C. ERα was detected using clone SP1 (Ready-to-Use, 32 minutes at 36°C; Ventana Medical Systems USA), PR using clone 1E2 (Ready-to-Use, 32 minutes at 36°C; Ventana Medical Systems, USA), AR using clone SP107 (1:300 dilution, 32 minutes at 37°C; Spring Bioscience, USA) and GR using clone D6H2L (1:600 dilution, one hour at 37°C; Cell Signalling Technologies, UK). To reduce background signal for the PR staining, after the primary antibody incubation step slides were incubated with normal antibody diluent (ABB999; Immunologic, Netherlands) for 24 minutes. ERα and PR were detected using the UltraView Universal DAB Detection Kit (Ventana Medical Systems, USA), while detection for AR was visualized using the OptiView DAB Detection Kit (Ventana Medical Systems, USA) and GR bound antibody was visualized using Anti-Rabbit HQ (Ventana Medical systems, USA) for 12 minutes at 37°C followed by Anti-HQ HRP (Ventana Medical systems, USA) for 12 minutes at 37°C, followed by the ChromoMap DAB detection kit (Ventana Medical Systems, USA).

### Rapid Immunoprecipitation of endogenous proteins (RIME)

Following hormone deprivation and treatment with hydrocortisone (2,75 µM) for two hours, RIME experiments were performed as previously described^7^. The following antibodies were used: anti-GR (12041, Cell Signalling Technology), anti-p57 (sc-56341, Santa Cruz Biotechnology), anti-rabbit IgG (sc-2027, Santa Cruz Biotechnology), and anti-mouse IgG (sc-2025, Santa Cruz Biotechnology).

Tryptic digestion of bead-bound proteins was performed as described previously^8^. LC-MS/MS analysis of the tryptic digests was performed on an Orbitrap Fusion Tribrid mass spectrometer equipped with a Proxeon nLC1000 system (Thermo Scientific) using the same settings, with the exception that the samples were eluted from the analytical column in a 90-min linear gradient.

Raw data were analysed by Proteome Discoverer (PD) (version 2.3.0.523, Thermo Scientific) using standard settings. MS/MS data were searched against the Swissprot database (released 2018_06) using Mascot (version 2.6.1, Matrix Science, UK) with Homo sapiens as taxonomy filter (20381 entries) for the GR-RIME experiment, whereas Sequest HT was used for the p57-RIME experiment. The maximum allowed precursor mass tolerance was 50 ppm and 0.6 Da for fragment ion masses. Trypsin was chosen as cleavage specificity allowing two missed cleavages. Carbamidomethylation (C) was set as a fixed modification, while oxidation (M) and deamidation (NQ) were used as variable modifications. False discovery rates for peptide and protein identification were set to 1% and as additional filter Mascot peptide ion score>20 or Sequest HT XCorr>1 was set. The PD output file containing the abundances was loaded into Perseus (version 1.6.1.3) [02] LFQ intensities were Log2-transformed and the proteins were filtered for at least 66% valid values. Missing values were replaced by imputation based on the standard settings of Perseus, i.e. a normal distribution using a width of 0,3 and a downshift of 1,8. Differentially expressed proteins were determined using a t-test (minimal threshold: FDR: 5% and S0: 0,1 for the p57-RIME experiment, for the GR-RIME experiment was the threshold–LOG(p-value) ≥ 2 and [x/y] ≥ 2,5 | [x/y] ≤ −2,5).). The comparison between the pooled cell lines of the p57 RIME experiment was IgG corrected.

### Immunofluorescence and quantification

After hormone deprivation cells were treated with 2,5 µM hydrocortisone or left untreated for 8 hours. Cells were washed and fixed in 2% paraformaldehyde for 10 minutes at room temperature. Subsequently, cells were permeabilized in 0.5% Triton-PBS for 10 minutes. After blocking for 60 minutes with blocking solution (1% BSA in PBS), samples were incubated for two hours with antibodies against GR (1:50), p57 (1:50) at room temperature. Following that, samples were incubated with secondary antibodies: Alexa 488 (A11001, Thermo Fisher Scientific) (1:1000) and Alexa 647 (A21244, Thermo Fisher Scientific) (1:1000) Finally, samples were counterstained with 4′,6-diamidino-2-phenylindole (DAPI) and analysed by either laser confocal microscopy (SP5, Leica) or screening fluorescent microscope (TIRF, Leica).

For single cell analysis, images were analysed in FIJI^9^. p57-positive cells were quantified in a fully automatic, unbiased manner, with a custom-made ImageJ macro script. For every image the DAPI channel was used to segment cell nuclei into ROIs as follows. After rolling ball background subtraction (40-micron radius) and a median filter (1,5-micron radius) local thresholding was applied (‘Mean’ method, 8-micron radius, with 4 times the standard deviation of the background as parameter), followed by a distance transform watershed operation to separate touching nuclei. The mean p57 signal was then measured inside the obtained ROIs. Cells were considered to be positive (negative) if this mean value was higher (lower) than a certain threshold, determined using untreated control samples. Resulting images with filled ROIs were overlaid with the original data for visual inspection.

### Drug screen

Before the start of the screen H1944 cells were cultured in medium without or with glucocorticoids (hydrocortisone, 2,75 µM) for two days. Using the Multidrop Combi (Thermo Fisher Scientific), untreated H1944 cells were seeded into 384-well plates either at low (1000 cells) or high (2500 cells) confluency, while the pre-treated H1944 cells were seeded at high (4500 cells) confluency. After 24 hours, the NKI compound collection of purchased drugs (Selleck GPCR, Kinase, Apoptosis, Phospatase, Epigenetic, LOPAC, and NCI oncology) was added. This library was stored and handled as recommended by the manufacturer. Compounds from the master plate were diluted in daughter plates containing complete RPMI-1640 medium, using the MICROLAB STAR liquid handling workstation (Hamilton). From the daughter plates, the diluted compounds were transferred into 384-well assay plates, in triplicate, with final concentrations of 1 μM, and 5 μM. In addition, Positive (1 µM Phenylarsine oxide) and negative (0,1% DMSO) controls were added alternately to wells in column 2 and 23 of each assay plate. After 6 days, viability was measured using CellTiter-Blue assay (G8081/2, Promega) following the protocol of the manufacturer. The CTB data was normalized per plate using the normalized percentage inhibition (NPI) method. NPI sets the mean of the positive control value to 0 and mean of the negative control to 1. When comparing GC-pre-treated vs vehicle and GC-co-treatment vs vehicle, the mean over the three replicates for each condition was calculated and then the vehicle mean was subtracted from the treated condition mean, producing the differential survival value. Using the replicate values of both conditions a two-sided t-test was performed. Afterwards the *p-values* were corrected for multiple testing using the Benjamini-Hochberg method^10^. All calculations were done in R.

### Phosphoproteomic analysis

After hormone depravation cells were treated with 2,75 uM hydrocortisone or left untreated (Veh) for 48 hours. For protein digestion, frozen cell pellets were lysed in boiling Guanidine (GuHCl) lysis buffer as previously described^11^. Protein concentration was determined with a Pierce Coomassie (Bradford) Protein Assay Kit (Thermo Scientific), according to the manufacturer’s instructions. Aliquots corresponding to 1,1 mg of protein were digested with Lys-C (Wako) for 2 hours at 37°C, enzyme/substrate ratio 1:100. The mixture was then diluted to 2 M GuHCl and digested overnight at 37°C with trypsin (Sigma-Aldrich) in enzyme/substrate ratio 1:100. Digestion was quenched by the addition of TFA (final concentration 1%), after which the peptides were desalted on a Sep-Pak C18 cartridge (Waters, Massachusetts, USA). From the eluates, aliquots were collected for proteome analysis, the remainder being reserved for phosphoproteome analysis. Samples were vacuum dried and stored at −80°C until LC-MS/MS analysis or phosphopeptide enrichment.

Phosphorylated peptides were enriched from 1 mg total peptides using High-Select Fe-NTA Phosphopeptide Enrichment Kit (Thermo Scientific), according to the manufacturer’s instructions, with the exception that the dried eluates were reconstituted in 15 μl of 2% formic acid.

Prior to mass spectrometry analysis, the peptides used for proteome analysis were reconstituted in 2% formic acid. Peptide mixtures were analysed by nanoLC-MS/MS on an Q Exactive HF-X Hybrid Quadrupole-Orbitrap Mass Spectrometer equipped with an EASY-NLC 1200 system (Thermo Scientific). Samples were directly loaded onto the analytical column (ReproSil-Pur 120 C18-AQ, 1,9 μm, 75 μm × 500 mm, packed in-house). Solvent A was 0,1% formic acid/water and solvent B was 0,1% formic acid/80% acetonitrile. Samples were eluted from the analytical column at a constant flow of 250 nl/min. For single-run proteome analysis, a 4h gradient was employed containing a linear increase from 7% to 30% solvent B, followed by a 15-minute wash, whereas for single-run phosphoproteome analysis a 2h linear gradient (from 4% to 22% solvent B, followed by a 15-minute wash) was used.

Proteome data (RAW files) were analysed by PD (version 2.3.0.523, Thermo Scientific) using standard settings. MS/MS data were searched against the human Swissprot database (20417 entries, release 2019_02) using Sequest HT. The maximum allowed precursor mass tolerance was 50 ppm and 0,06 Da for fragment ion masses. Trypsin was chosen as cleavage specificity allowing two missed cleavages. Carbamidomethylation (C) was set as a fixed modification, while oxidation (M) and deamidation (NQ) were used as variable modifications. False discovery rates for peptide and protein identification were set to 1% and as additional filter Sequest HT XCorr>1 was set. The PD output file containing the abundances was loaded into Perseus (version 1.6.1.3) [02]. LFQ intensities were Log2-transformed and the proteins were filtered for at least two out of three valid values in one condition. Missing values were replaced by imputation based on the standard settings of Perseus, i.e. a normal distribution using a width of 0,3 and a downshift of 1,8. Differentially expressed proteins were determined using a t-test (threshold: p ≤ 0,05 and [x/y] ≥ 1,5 | [x/y] ≤ - 1,5).

Phosphoproteome data (RAW files) were analysed by MaxQuant (version 1.6.1.0) using standard settings^12^. MS/MS data were searched against the human Swissprot database (20,417 entries, release 2019_02) complemented with a list of common contaminants and concatenated with the reversed version of all sequences. The maximum allowed mass tolerance was 4,5 ppm in the main search and 20 ppm for fragment ion masses. False discovery rates for peptide and protein identification were set to 1%. Trypsin/P was chosen as cleavage specificity allowing two missed cleavages. Carbamidomethylation (C) was set as a fixed modification, while oxidation (M), deamidation (NQ) and phosphorylation (S,T,Y) were used as variable modifications. LFQ intensities were Log2-transformed in Perseus (version 1.6.5.0), after which the phosphosites were filtered for at least two valid values (out of 3 total) in both conditions. Missing values were replaced by imputation based a normal distribution using a width of 0,3 and a downshift of 1,8. Differentially regulated phosphosites were determined using a t-test (threshold: p ≤ 0,05 and [x/y] ≥ 1,5 | [x/y] ≤ −1,5. These differential phosphosites were combined with on/off (3 out of 3 total present/missing) phosphosites. For the cytoscape analysis the app PhosphoPath was used. Data was loaded into the PhosphoPath plug-in and processed as described in the manual^13^.

### Time-Lapse Microscopy

For doubling time experiments, a Lionheart FX automated microscope was used. Cells (∼10 000 per well) were plated in a 96 well plate and sirDNA77 with or without 2,75 µM hydrocortisone was added one hour before imaging. Growth curves were generated with a time resolution of 4 hours for a total time span of 144 hours (microscope maintained at 37°C, 5% CO2 using a 4x lens and a Sony CCD, 1,25-megapixel camera with 2 times binning; BioTek). Quantification of cell number was performed by Gen5 software (BioTek). Doubling times were calculated using GraphPad Pism 6 software.

For mitotic entry experiments, cells (∼20 000 per well) were grown on Lab-Tek II chambered coverglass (Thermo Scientific). One hour before imaging, 2,75 uMf hydrocortisone was added per condition. Images were obtained every 15 minutes during 60 hours using a DeltaVision Elite (applied precision) maintained at 37°C equipped with a 10x PLAN Apo S lens (Olympus) and cooled CoolSnap CCD camera. Up to five images were acquired per well and 50 cells per experiment were evaluated. Image analysis was performed using ImageJ software (NIH). The percentage of mitotic entry was determined following cells from the start of the movie until they divided up to 60 hours.

### 4C analysis

4C was performed as previously described^14^ with minor modifications^15^. 4C libraries were sequenced on a MiSeq and were analysed with a custom 4C mapping pipeline (https://github.com/deWitLab/4C_mapping). 4C ligation data were mapped to hg19. Normalization and downstream analysis were done using peakC^16^.

### Breast cancer genetic models

Activity of GR in mouse mammary tumours was assessed in bulk tumour RNA sequencing data as previously described^17^. Briefly, RNA was extracted from tumours of using TRIzol reagent (Thermo Fisher Scientific). Tumours from the following mouse models were analysed: *Keratin 14 (K14)-cre;Cdh1^F/F^;Trp53^F/F^*, *K14-cre;Brca1^F/F^;Trp53^F/F^, K14-cre;Brca1^F/F^;Trp53^F/F^;Col1a1^invCAG-IRES-Luc/+^* (*K14-cre;Brca1^F/F^;Trp53^F/F^;Col1a1*), *K14-cre;Brca1^F/F^;Trp53^F/F^;Col1a1^invCAG-Met-IRES-Luc/+^* (*K14-cre;Brca1^F/F^;Trp53^F/F^;Met*), *Whey acidic protein (Wap)-cre;Brca1^F/F^;Trp53^F/F^*, *Wap-cre;Brca1^F/F^;Trp53^F/F^;Col1a1^invCAG-Myc-IRES-Luc/+^* (*Wap-cre;Brca1^F/+^;Trp53^F/F^;Myc*), *Wap-cre;Brca1^F/F^;Trp53^F/F^;Col1a1^invCAG-Myb2-IRES-Luc/+^* (*Wap-cre;Brca1^F/F^;Trp53^F/F^;Myb2*), *Wap-cre;Cdh1^F/F^;Col1a1^invCAG-Akt1E17K-IRES-Luc/+^* (*Wap-cre;Cdh1^F/F^;Akt1^E17K^*), *Wap-cre;Cdh1^F/+^;Col1a1^invCAG-Akt1E17K-IRES-Luc/+^* (*Wap-_cre;Cdh1_F/+_;Akt1_E17K*_), *Wap-cre;Cdh1*_*F/F_;Col1a1_invCAG-Pik3caE545K-IRES-Luc/+* _(*Wap-*_ *cre;Cdh1^F/F^;Pik3ca^E545K^*) and *Wap-cre;Map3k1^F/F^;Pten^F/F^*. Samples were single-end sequenced for 51 or 65 base pairs on the Illumina Hiseq2000/Hiseq2500 Machine. The reads were aligned against the mouse transcriptome (mm10) using Tophat2 (Tophat version 2.1.0, Bowtie version 1.0.0). Tophat was guided using a reference genome as well as a reference transcriptome. Differential expression analysis was performed using the R package edgeR in combination with the voom method, using raw read counts as input. Library size normalization was performed during differential expression analysis within the voom function. Genes with *p*<0.05 were considered as differentially expressed. GR activity score was subsequently analysed by assessing the differential expression of GR target genes^18^ between *Trp53^F/F^* and *Trp53^+/+^* mouse tumours.

### Copy number profiling

Copy number profiles were generated using ChIP-seq input samples. The reads were combined in 20kb bins across the genome, and data analysed as previously described^19^.

### GR-ChIP seq and Hi-C time course analysis

Chromatin loops spanning either FOXO1 or IRS2 loci in A549 previously identified using Hi-C^20^ were analysed. Dynamic loops showing a significant increase or decrease in chromatin interaction frequency after 1, 4, 8 and 12 hours of dexamethasone exposure were reported. GR and c-Jun (AP-1) binding sites identified by ChIP-seq in a 12 hours dexamethasone timecourse^21^ that significantly gained signal and that overlapped a 10kb anchor of a dynamic loop were interrogated. The changes induced by dexamethasone for GR and c-Jun ChIP-seq signal and chromatin interaction counts were fitted into generalized linear models (GLMs) using edgeR^22^ and a likelihood ratio test was performed to identified significant hits (FDR≤0,05) as previously described. Log2 fold-change values were calculated for each dexamethasone timepoint over the absence of dexamethasone.

### Statistical analysis

Statistical analysis was performed using Prism (GraphPad, San Diego, CA). Normality was tested using D’Agostino-Person and Shapiro-Wilk test, if data was normally distributed Student’s t-test (unpaired or Welch’s t test when variance is unequal) was applied, whereas in case of deviation from normal distribution Mann–Whitney U test was used. For comparison of multiple groups Kruskal-Wallis test was used. Technique-specific statistical tests are described within their corresponding method subsection.

---

**Figure S1.**
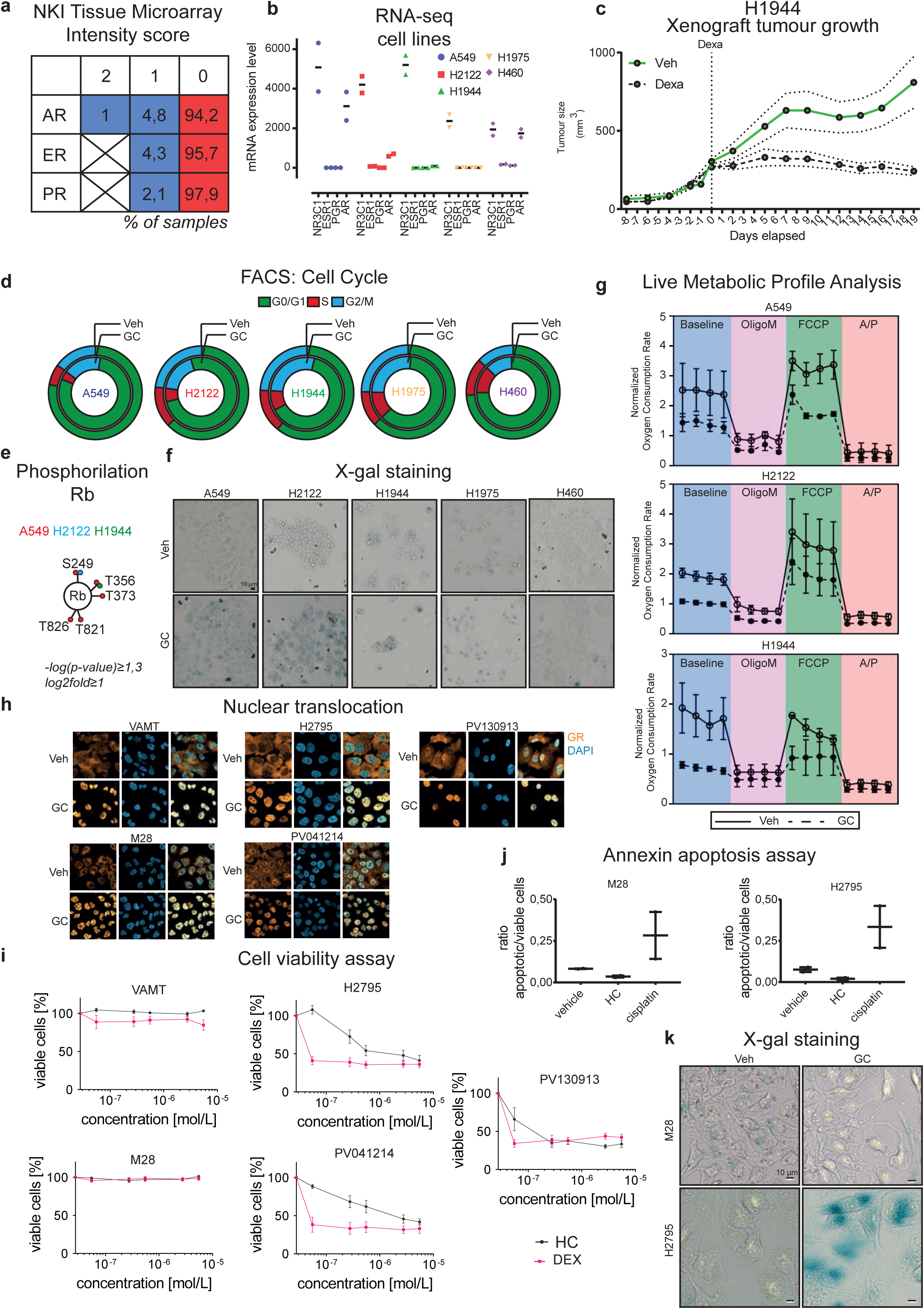
**a,** Immunohistochemistry of steroid hormone receptors in lung cancer. AR (193), ER (47), PR (47) (top left) staining scores with percentages. **b,** mRNA expression level of steroid hormone receptors in lung cancer cell lines. n=2. **c,** Xenograft tumour growth in NOD/SCID mice. n≥7. **d,** Cell cycle distribution by phase of untreated (Veh) or glucocorticoid treated (GC) cells. **e,** Phosphosites of pocket protein Rb found to have significant increase in phosphorylation upon GC treatment in responders. n=3. **f,** Representative X-gal staining images of untreated (Veh) or glucocorticoid treated (GC) lung cell lines. n=3. **g,** Per DNA content normalized oxygen consumption rate. For each condition four timepoints were measured. OligoM = Oligomycin; FCCP = arbonyl cyanide-4-(trifluoromethoxy)phenylhydrazone; A/R = Antimycin A/Rotenone. Vehicle (Veh; Full lines) and glucocorticoid (GC; dotted lines) treatment was used. n≥2, five technical replicates each. Bars depict mean ±SEM. **h,** Representative immunofluorescence images showing expression and localization of GR (orange), treated with glucocorticoids (GC) or control (Veh), using DAPI as nuclear staining (light blue) in mesothelioma cell lines. **i,** Cell viability assays survival of mesothelioma cell lines treated with increasing concentrations of glucocorticoids (HC = hydrocortisone or DEX = dexamethasone). n=3. **j,** Ratio of apoptotic to viable cells in two mesothelioma cell lines untreated (Veh), or treated with glucocorticoids (GC) or cisplatin. n=2. **k,** Representative X-gal staining images of untreated (Veh) or glucocorticoid treated (GC) mesothelioma cell lines. n=3.

**Figure S2.**
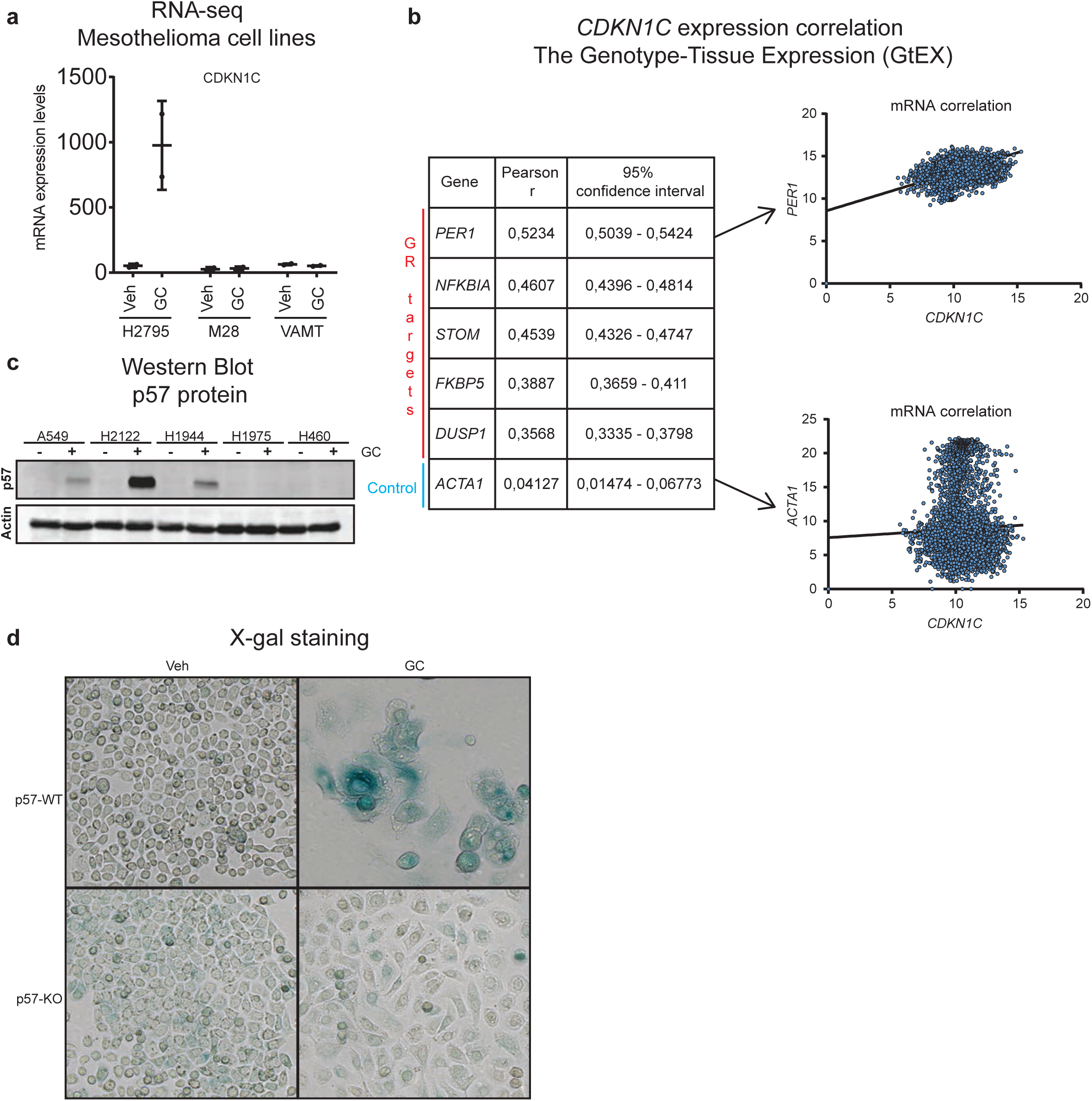
**a,** mRNA expression levels of *CDKN1C* in mesothelioma cell lines, untreated (Veh) or treated with glucocorticoids (GC). n=2. **b,** Expression correlation of GR target genes with *CDKN1C* across different human tissues (excluding blood, brain, and skin). **c,** Representative western blot showing p57 expression, with actin as loading control. n=2**. d,** Representative X-gal staining images of untreated (Veh) or glucocorticoid treated (GC) cells, H2122 p57-WT and p57-KO. n=2

**Figure S3.**
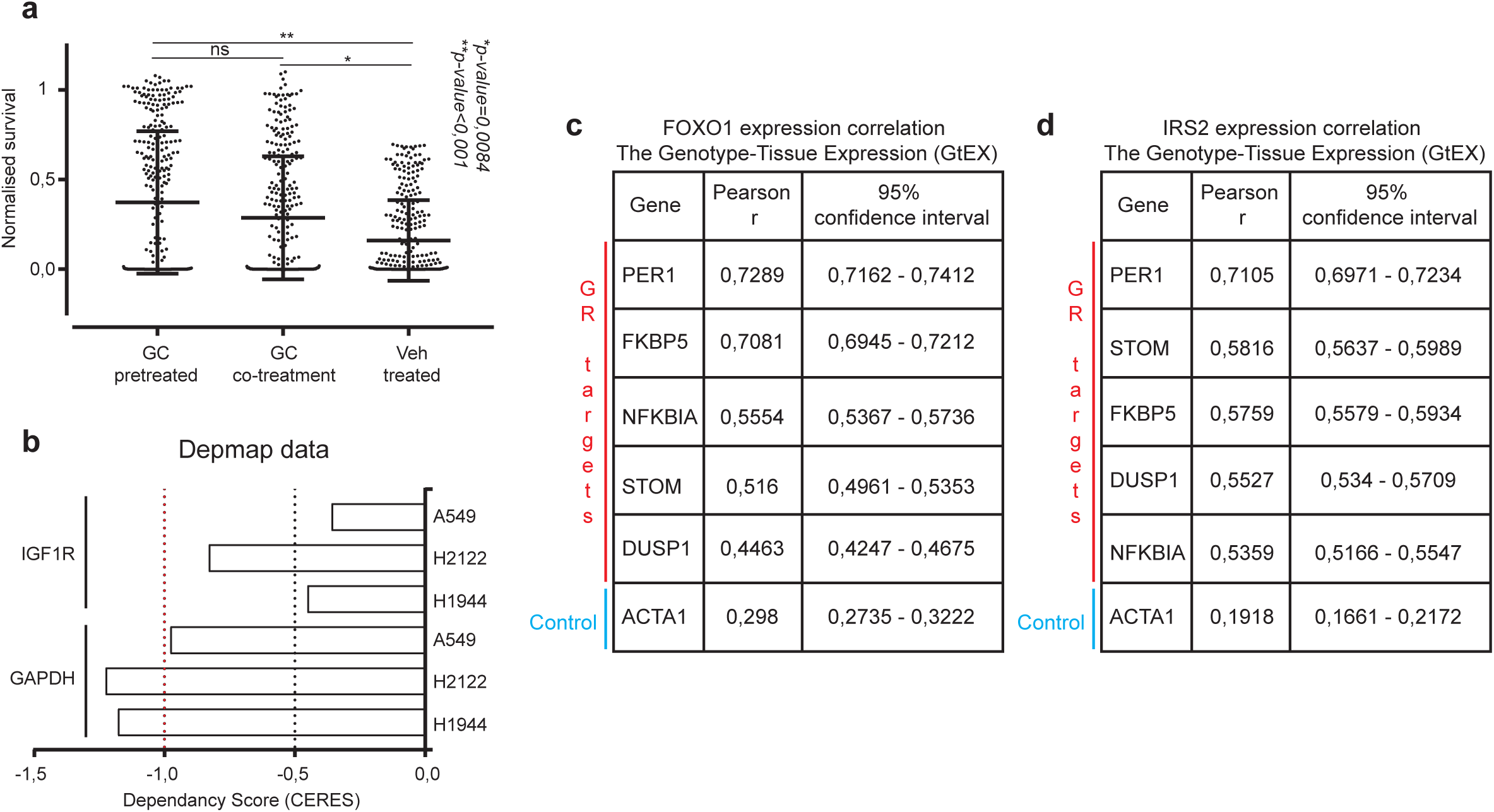
**a,** Normalised survival for compounds giving significantly reduced viability of vehicle (Veh; Normalised survival ≤0,7) treated H1944 cells and their effects on the other two arms in the screen. Bars depict mean ±SD. **b,** Dependency scores (CERES) for *IGF1R* and *GAPDH* for A549, H2122, and H1944 cell lines. **c,** Expression correlation of GR target genes with *FOXO1* (left) and *IRS2* (right) across different human tissues (excluding blood, bone marrow, brain, and skin).

**Figure S4.**
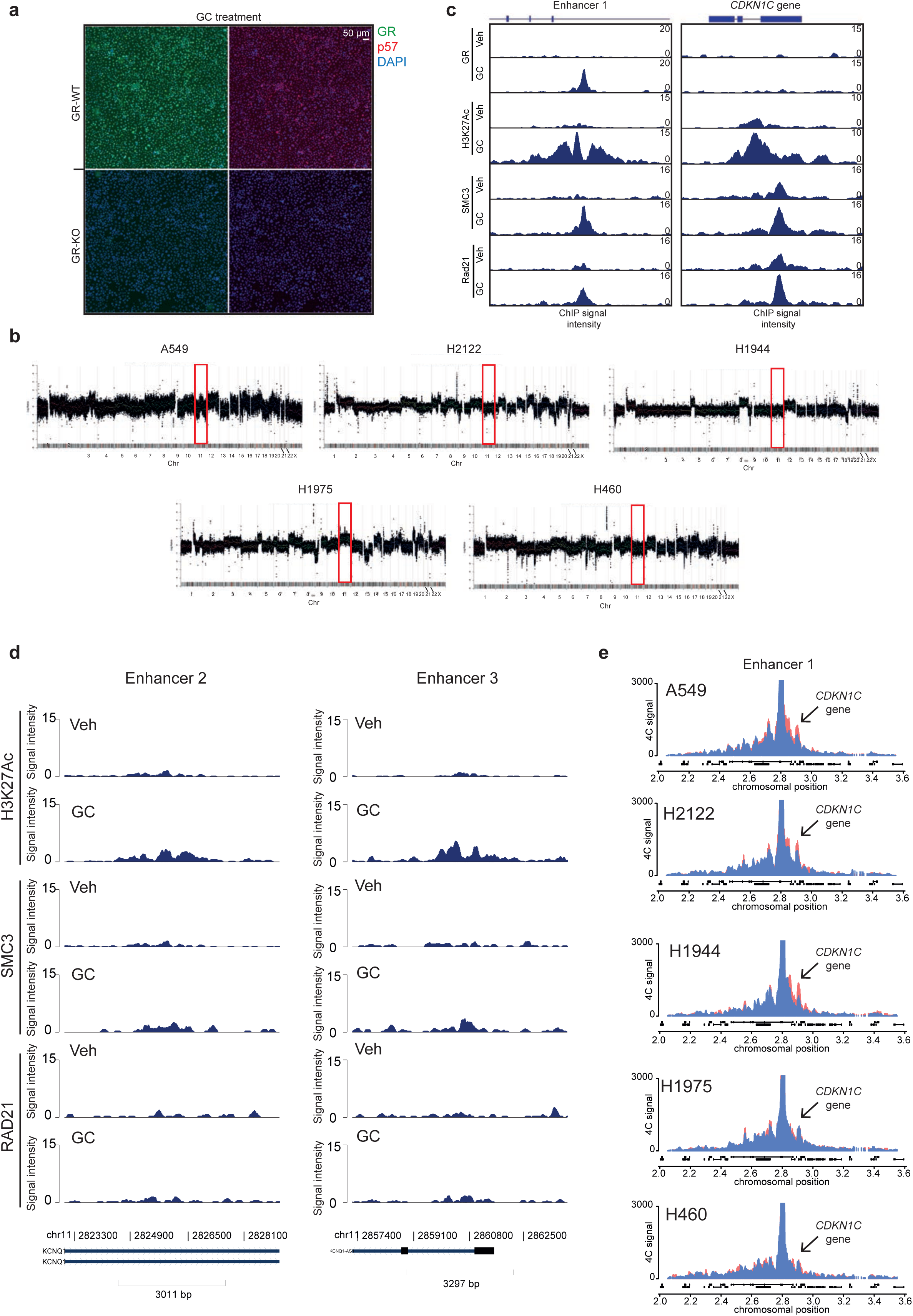
**a,** Representative immunofluorescence images showing expression and localization of GR (green) and p57 (red) in GR-WT and GR-KO H2122 cell lines. DAPI is used as nuclear staining (blue). Scale bar, 50 μm. n=2. **b,** Genome-wide copy number status (Log Ratio) per probe. **c,** ChIP sequencing data (ENCODE 01HG007900) showing peaks for GR, H3K27Ac, SMC3, and Rad21 at Enhancer 1, and CDKN1C gene in A549 cell line, not treated (Veh) or treated (GC). **d,** ChIP sequencing data (ENCODE 01HG007900) showing peaks for GR, H3K27Ac, SMC3, and Rad21 at Enhancer 2 and 3, in A549 cell line, not treated (Veh) or treated (GC). **e,** 4C signal across the region surrounding Enhancer 1 on chromosome 11, under Vehicle (Veh; blue) and glucocorticoid (GC; red) treatment. Average signal of 2 biological replicates shown.

**Figure S5.**
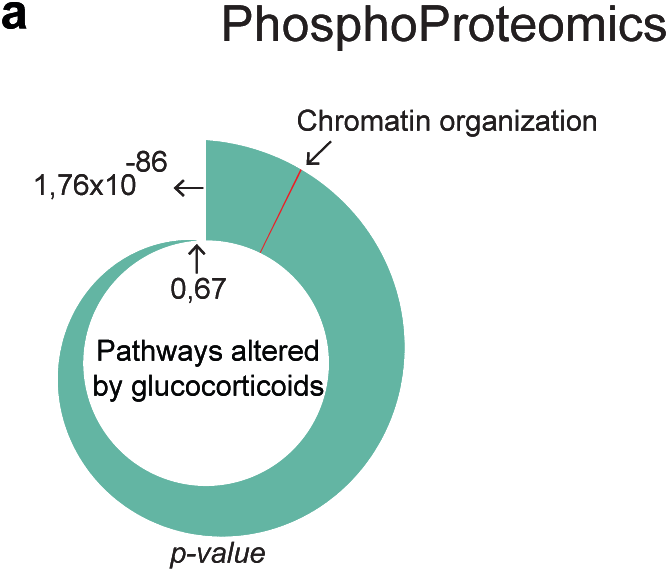
**a,** PhosphoPath pathway analysis identified “Chromatin Organization” pathway enrichment in GC treated cells (A549, H2122, and H1944). n=3.

**Figure S6.**
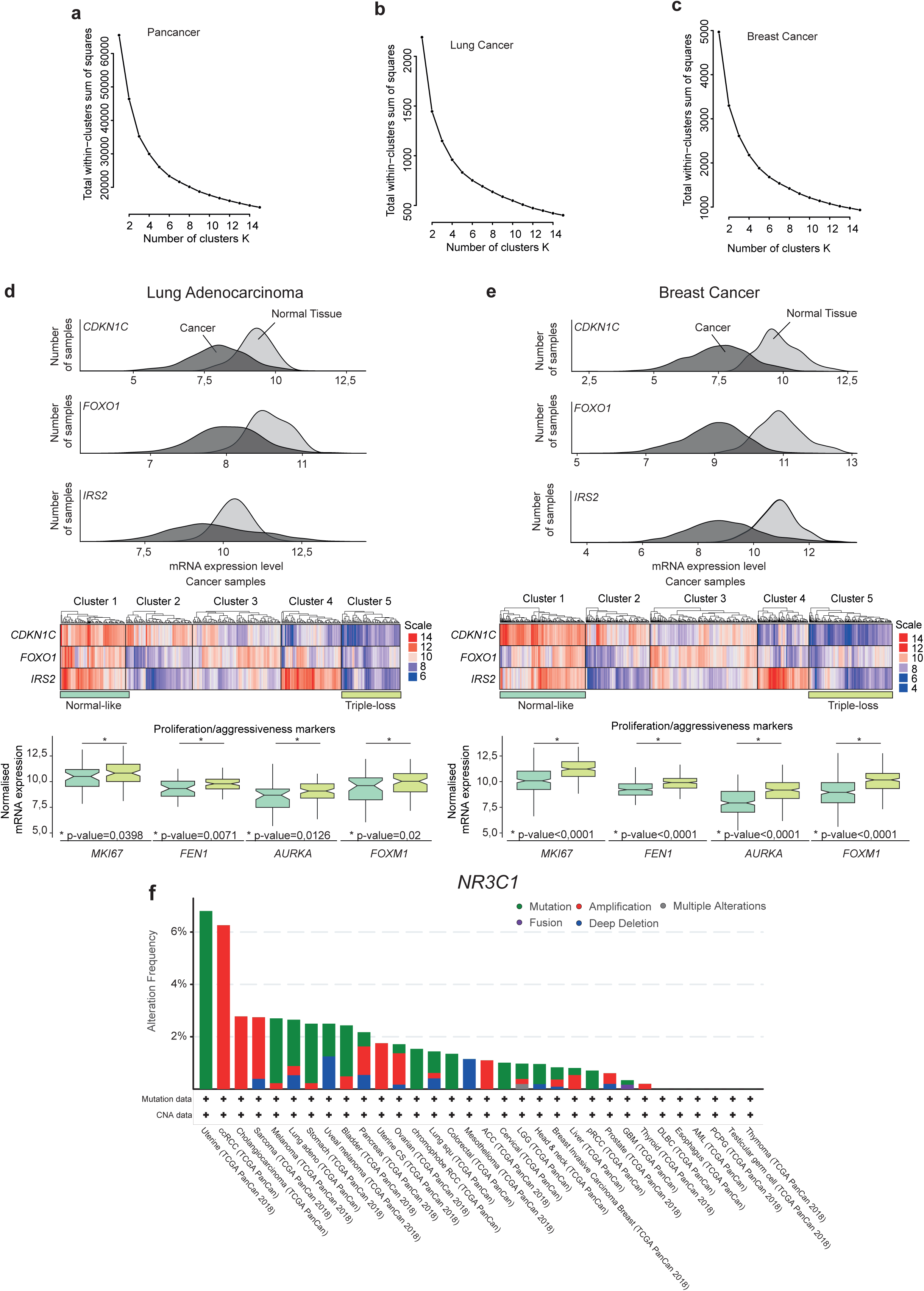
**a-c,** Elbow plot method depicts total within−clusters sum of squares as a function of number of clusters (K) **d-e,** Ridged plots showing normalised mRNA expression of *CDKN1C*, *FOXO1*, and *IRS2* across normal (light grey) and tumour (dark grey) specimens (top). Unsupervised clustering (k_means_=5) of cancer samples (middle). Normalised mRNA expression of proliferation/aggressiveness markers (*MKI67*, *FEN1*, *AURKA*, and *FOXM1*) between ‘normal-like’ and ‘triple-loss’ cluster (bottom). **f,** Alterations in *NR3C1* gene (coding for GR) across various cancer types.

## Notes

#### Summary of Updates

Author Secuk Y. corrected to Selcuk, L was missing. Author Raoderick B. corrected to Roderick B, a was inserted by accident.

